# Profiling genome-wide methylation in two maples: fine-scale approaches to detection with nanopore technology

**DOI:** 10.1101/2022.08.02.502577

**Authors:** Susan L. McEvoy, Patrick G. S. Grady, Nicole Pauloski, Rachel J. O’Neill, Jill L. Wegrzyn

## Abstract

DNA methylation is critical to the regulation of transposable elements and gene expression, and can play an important role in the adaptation of stress response mechanisms in plants. Traditional methods of methylation quantification rely on bisulfite conversion that can compromise accuracy. Recent advances in long-read sequencing technologies allow for methylation detection in real time. The associated algorithms that interpret these modifications have evolved from strictly statistical approaches to Hidden Markov Models and, recently, deep learning approaches. Much of the existing software focuses on methylation in the CG context, but methylation in other contexts is important to quantify, as it is extensively leveraged in plants. Here, we present methylation profiles for two maple species across the full range of 5mC sequence contexts using Oxford Nanopore Technologies (ONT) long-reads. Hybrid and reference-guided assemblies were generated for two new *Acer* accessions: *Acer negundo* (65x ONT and 111X Illumina) and *Acer saccharum* (93x ONT and 148X Illumina). The ONT reads generated for these assemblies were re-basecalled, and methylation detection was conducted in a custom pipeline with the published *Acer* references (PacBio assemblies) and hybrid assemblies reported herein to generate four epigenomes. Examination of the transposable element landscape revealed the dominance of *LTR Copia* elements and patterns of methylation associated with different classes of TEs. Methylation distributions were examined at high resolution across gene and repeat density and described within the broader angiosperm context, and more narrowly in the context of gene family dynamics and candidate nutrient stress genes.

## INTRODUCTION

The processes shaping plant development and growth are regulated by epigenetic modifications that impact gene expression, genomic stability, and plasticity (Kumar and Mohapatra 2021). Plants leverage methylation in sequence contexts beyond the CG dinucleotide (CHG and CHH, where H = C, A, or T), and these modifications are primarily regulated by transposable elements (TEs), which represent a significant portion of most plant genomes. Epigenetic modifications introduced through mobile elements contribute to genetic variation that is associated with biotic and abiotic stress adaptations (Ritter and Niederhuth 2021).

Methylation in model plants has been studied from its initiation in embryos in *Arabidopsis thaliana* (Jullien et al. 2012), to the accumulation of methylation variation in the independent branches of a single *Populus trichocarpa* individual (Hofmeister et al. 2020). In *Arabidopsis*, the genome-wide regulatory effects of methylation were demonstrated by the knockout of all five methyltransferases, impacting cell fate throughout the plant (He et al. 2022). Also in *Arabidopsis*, a heritable epiallele associated with climate was found to control leaf senescence, providing an example of local climate adaptation (He et al. 2018). In *P. trichocarpa*, epigenetic modifications were associated with changes in the circadian cycle (Liang et al. 2019). Increasingly, non-model plant systems have been investigated, including for methylation responsible for leaf shape and photosynthetic traits in *Populus simonii* (Ci et al. 2016) and salinity-induced methylation in mangroves (Miryeganeh et al. 2022).

Until recently, methylation was primarily investigated through treatments such as whole-genome bisulfite conversion (WGBS). This technique is prone to degradation of DNA, incomplete conversion, and amplification bias (Gouil and Keniry 2019). Since WGBS libraries are primarily short-read sequenced, interpretation also suffers from poor resolution in repetitive regions. Long-read technologies, including Pacific Biosciences’ (PacBio’s) single-molecule real-time (SMRT) sequencing, and nanopore sequencing by Oxford Nanopore Technologies (ONT), can detect methylation. SMRT sequencing can detect 5mC modifications based on polymerase dynamics at very high coverage, as well as methods that rely on bisulfite conversion for standard coverage. In comparison, nanopore sequencing can directly detect DNA or RNA modifications through a voltage-measured pore, enabling real-time, single-molecule sensitivity (Liu et al. 2021).

Being sessile, plants rely on heritable methylation as an evolutionary strategy, and this is particularly important in long-lived tree species. DNA methylation is known to have a critical role in the silencing of TEs, but environmental stress can reduce this activity and result in TE bursts (Cavrak et al. 2014). Epigenetic mechanisms related to transposition have been associated with responses to drought, temperature, and nutrient stress (Fan, Peng, and Zhang 2022). In the context of maples, sugar maple (*A. saccharum*) is susceptible to calcium deficiency, and this has led to a significant decline in natural populations (Bishop et al. 2015). The first comparative genomics study on North American maples identified candidate genes from the analysis of expression in the aluminum-and calcium-amended plots at the Hubbard Brook Experimental Forest (McEvoy et al. 2021). The interplay between TEs and gene expression is complex, but many forest tree species would benefit from a deeper examination to fully understand their adaptive potential for new and ongoing threats.

This study extends the previous work on *A. negundo* and *A. saccharum* (McEvoy et al. 2021), as well as that of Sork et al. (2021) and Niederhuth et al. (2016) in comparative plant methylomics. Employing ONT sequencing from two previously unstudied *Ace*r individuals, we completed genome assembly and annotation and detected methylation sites genome-wide. We customized methylation calling methods, and in doing so, generated comparative methylation profiles focusing on transposable elements and nutrient stress candidate genes.

## METHODS

### Sequencing

Leaves from two maple individuals, *A. negundo* (Accession 253-2013*B) and *A. saccharum* (1353-84*A), were shipped on dry ice from the Arnold Arboretum of Harvard University. High molecular weight (HMW) DNA was extracted from both samples using the ONT protocol for *Arabidopsis* leaves (Vaillancourt and Robin Buell 2019). The resulting gDNA was checked for quality control via Thermo Scientific Nanodrop and then Agilent TapeStation (Agilent Technologies, Santa Clara, CA). Libraries were prepared with the ONT Genomic DNA by Ligation protocol with additional Covaris shearing to improve coverage (Oxford Nanopore Technologies, Oxford, UK). A single ONT Flongle sequencing run was conducted to evaluate the library quality prior to the PromethION run (*A. negundo*: 15 K reads, N50 16 Kb in 23 hours; *A. saccharum*: 39 K reads, N50 14 Kb after 21 hours). Two PromethION runs (one per individual) followed, using flow cell type FLO-PRO002, kit SQK-LSK110, and Guppy v4.0.11 (Oxford Nanopore Technologies 2020a) with the high-accuracy basecalling model and read filtering minimum qscore of 7. To reduce the error rate, particularly for methylation calling, the resulting FAST5s were re-basecalled with Guppy v5.0.16 (GPU) and the latest ONT super-accuracy model. The same samples were short-read sequenced in a single Illumina NovaSeq 6000 SP v1.5 300 cycle run. TruSeq DNA Nano with Covaris shearing was used for Illumina library preparation in advance (Illumina Inc., San Diego, CA, USA).

### Assembly and annotation

ONT long-reads were filtered for archaea, bacteria, fungi, and virus contaminants via Centrifuge v1.0.4-beta (Kim et al. 2016) as well as length (5 Kb minimum). Illumina short reads were quality-controlled with FASTP v0.22.0 (Chen et al. 2018). The filtered ONT reads were combined with raw Illumina reads for hybrid assembly with MaSuRCA v4.0.3 (Zimin et al. 2017). This was followed by short-read polishing with Pilon v1.24 (Walker et al. 2014) using the trimmed short-reads that were aligned with Bowtie v2.3.5.1 (Langmead et al. 2009). Scaffolding was performed with RagTag v2.1.0 (Alonge et al. 2021) using the original chromosome-scale reference genomes for the same species (McEvoy et al. 2021).

Repeats were identified with RepeatModeler v2.0.1 (Flynn et al. 2020) and combined with the LTR reference library InpactorDB non-redundant v5 (Orozco-Arias et al. 2021) for softmasking with RepeatMasker v4.0.6 (Smit, Hubley, and Green 2013–2015). ParseRM generated repeat summaries and estimates of abundance by divergence. BRAKER2 v.2.1.6 predicted genes with the previously published RNA-Seq leaf tissue library provided as evidence (Brůna et al. 2021) The RNA-Seq leaf tissue libraries represented 31.2 M read pairs for *A. negundo* and 30.6 M read pairs for *A. saccharum*. Gene models were filtered with gFACs v1.1.3 (Caballero and Wegrzyn 2019) with the following: unique genes only, mono-exonics missing a start or stop codon or valid protein domain, mult-iexonics missing both a start and stop codon, and genes with exons smaller than 6 bp. Functional annotation of the final gene space was conducted with EnTAP v0.10.7 using RefSeq Complete and Uniprot NCBI databases, along with Eggnog v4.1 for gene family assignment (O’Leary et al. 2016; Hart et al. 2018; UniProt Consortium 2019).

### Methylation detection

The METEORE pipeline was selected as the general approach to methylation detection based on its ability to generate reliable consensus results from multiple tools (Yuen et al. 2021). Two tools were selected: Nanopolish v0.13.2 (Loman, Quick, and Simpson 2015) and DeepSignal-Plant v0.1.4 (Ni et al. 2021). This pairing was chosen as it had favorable results in the Yuen et al. (2021) benchmarking study and the ability to inform beyond the CG-context. Nanopolish is a well-supported tool that detects methylation in the CG context. DeepSignal-Plant is the top-performing, most accessible tool trained with plant-based models to detect methylation in CHG and CHH contexts from nanopore sequencing (Ni et al. 2021).

ONT reads from the new individual were aligned to both the new and original genomes for methylation calling. Methods for Nanopolish and DeepSignal-Plant proceeded according to the documentation for each tool and the Snakemake workflows provided by METEORE (Fig. 1). Each tool was run to create two output formats: 1) the input format necessary for integration in the consensus, and 2) the standardized form of independent tool output, calculating per-site and per-strand frequencies, allowing for greater ease of interpretation across tools. This provided both tool-specific frequencies and a consensus for comparison.

**Figure 1.**
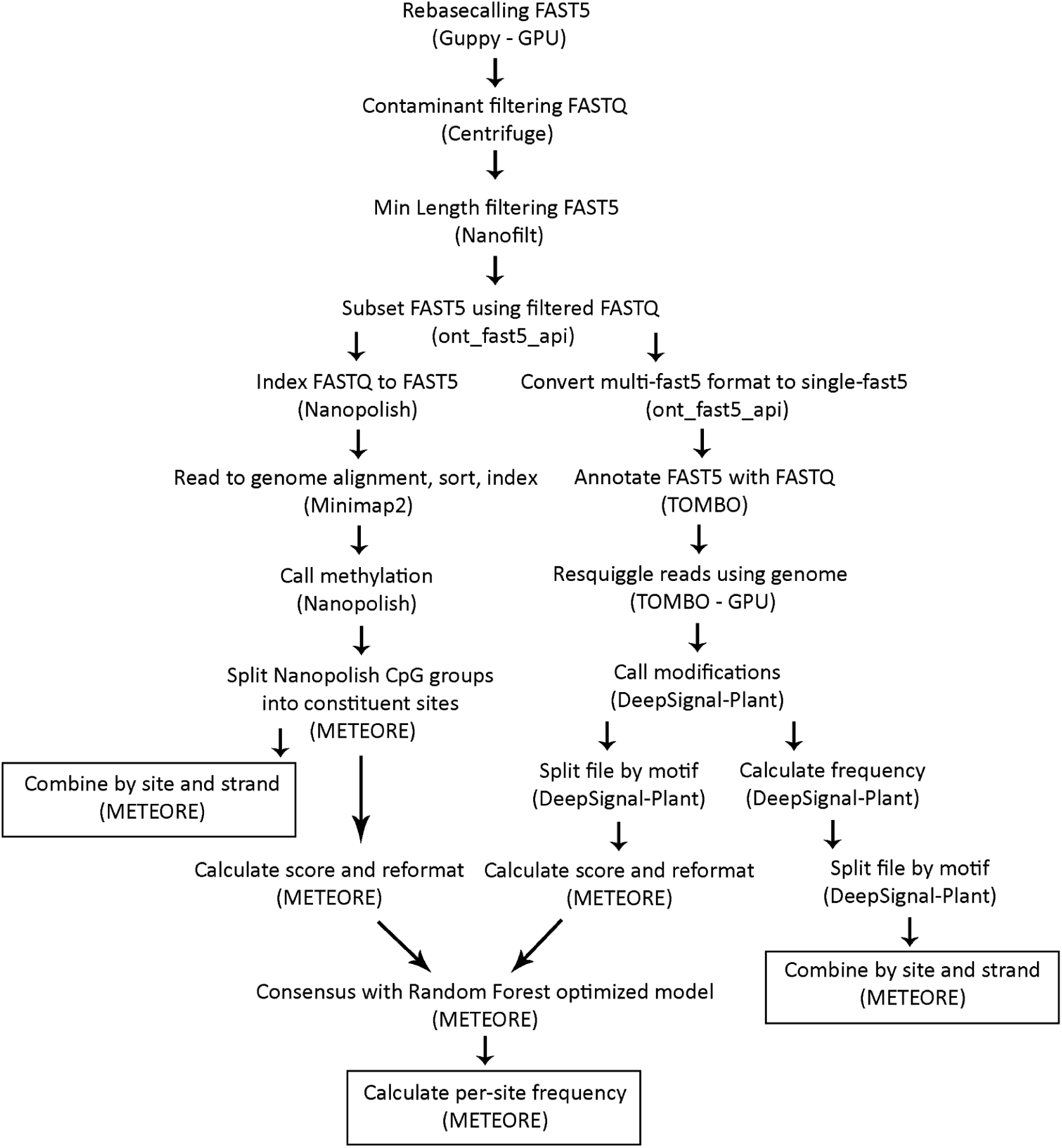
Stepwise method of methylation detection using the METEORE pipeline to create a consensus of DeepSignal-Plant and Nanopolish.

To begin, the rebasecalled and filtered FASTQ reads were used to filter the FAST5 using ont_fast5_api ‘fast5_subset.py’ v4.0.0 (Oxford Nanopore Technologies 2021). Nanopolish requires indexing of FASTQ to FAST5 in the default multi-fast5 format, and then alignment with minimap2 with parameters-ax map-ont (H. Li 2018). These alignments were run with minimap2 version 2.22-r1101 outside of the METEORE pipeline because the within-METEORE version provided by Snakemake was older. The next step was Nanopolish ’call_mods’, followed by METEORE scripts to convert the output of log-likelihood ratio values into a standardized format.

DeepSignal requires single-fast5 formatted files, so the multi-fast5 files were converted using ont_fast5_api ‘multi_to_single_fast5’. DeepSignal-Plant began with annotation of the FAST5s with FASTQ using Tombo ‘preprocess annotate_raw_with_fastqs’ v1.5.1 (Oxford Nanopore Technologies 2020b). This was followed by Tombo ‘resquiggle’ with the parameter--signal-align-parameters 4.2 4.2 2000 2500 20.0 40 750 2500 250 used for the original genomes only. It is possible to detect methylation on either reads or extracted features; reads were recommended, so this method was implemented. At this stage, results were split into separate files for CG, CHG, CHH, and CHH subcontexts by modifying METEORE split_freq_file_by_5mC_motif_sm.py. METEORE scripts were then used to standardize per-site and per-strand formats as described for Nanopolish. The CG file alone was formatted as input for the consensus script. Consensus predictions were generated using a Random Forest with provided models optimized at n-estimator = 3 and max_dep = 10. Results were analyzed with BEDTools v2.29.0 (Quinlan and Hall 2010) and karyoplotR (Gel and Serra 2017).

### Statistical analysis and visualization

BEDTools v2.29.0 (Quinlan and Hall 2010) was used to create 1 Mb windows and 100 Kb windows with 50 Kb overlaps across the genomes. It was then used to map methylation to these windows and calculate the frequency mean. BEDTools was also used with gene annotation files to intersect the gene regions and count methylated sites within the region. Plotting of chromosomal distributions was conducted with karyoplotR v1.21.3 (Gel and Serra 2017).

Pearson correlation was calculated with R Core Stats Package v4.2.0 (R Core Team 2013). Statistics and summaries were used to compare across reference genomes within *Acer* (McEvoy et al. 2021), as well as *Populus* (Hofmeister et al. 2020), *Quercus* (Sork et al. 2022), and the 34 angiosperms surveyed by Niederhuth (2016). Rapidly expanding and contracting families within *A. negundo* and *A. saccharum* were plotted along the distributions of methylation frequency and TE coverage. A total of 245 candidate genes associated with calcium response in *A. saccharum* were also investigated.

Scripts for all methods are available at https://gitlab.com/PlantGenomicsLab/acermethylation

## RESULTS AND DISCUSSION

### Sequencing

The ONT sequencing of *A. negundo* resulted in 48.5 Gb (115x coverage) in 4.29 M reads with a read N50 of 16.3 Kb. Re-basecalling resulted in an expected loss, resulting in 29.6 Gb (70x) in 2.55 M reads with an N50 of 16.2 Kb. After contaminant (56 K reads) and length filtering, 65x coverage with an N50 of 16.6 Kb remained. Illumina raw reads had 111x coverage as 46.6 Gb in 310 M reads. After trimming for adaptors, length, and quality, 104x coverage (146 M total read pairs) remained (<u>File S1</u>). Nanopore sequencing of *A. saccharum* resulted in 104.5 Gb in 9.8 M reads with an N50 of 15.1 Kb. Re-basecalling resulted in 56.7 Gb (99x coverage) in 5.2 M reads with an N50 of 15.2 Kb. Filtering (135 K contaminant reads) generated coverage of 93x with a read N50 of 15.5 Kb. Illumina sequencing (150bp PE) produced 148x coverage of 52.9 Gb bases in 563 M read pairs and trimming reduced this to 139x (526 M total read pairs).

### Assembly and annotation

After the initial draft assembly, *A. negundo* had a total length of 421 Mbp in 421 contigs and an N50 of 2.16 Mb. BUSCO embryophyta genes were 96.0% complete, with 2.9% of these in duplicate. Polishing only minimally reduced the length and N50 (<u>File S1</u>). Scaffolding with the original genome increased the length slightly, but it remained within 421 Mbp, and the N50 grew to 33 Mb. The assembly was in 32 scaffolds, with 13 chromosomes representing 99.7% of the assembled length. The genome size remained constant, though slightly smaller, than the published reference (442 Mbp) and closer to the kmer-based estimate of 319 Mbp (McEvoy et al. 2021). At 96%, the final complete assembly BUSCO scores remained similar to the draft, with duplicates dropping slightly to 2.8%, 0.7% fragmented, and 3.3% missing (Table 1, <u>File S1</u>). The lower duplicate value was an improvement from the original reference duplication of 3%, although the completed single copy dropped slightly. Structural annotation of the new genome identified 27,541 genes, of which 23,408 were functionally annotated by either similarity search or gene family assignment. The BUSCO score for annotated proteins was 92.1% complete, with 3% in duplicates (Table 1).

**Table 1.** Assembly and annotation statistics for new and original genomes

| Assembly | Genome |  |  | Genes |  |  |  |  |
| --- | --- | --- | --- | --- | --- | --- | --- | --- |
|  | Length | N50 | BUSCO* | Total | Mono-exonic | Average length | Avg. CDS length | BUSCO* |
| **A. negundo v2** | 421 Mbp | 33 Mb | 96.0 (2.8) | 27,541 | 3474 | 3098 | 1122 | 92.1 (3.0) |
| **A. negundo v1** | 442 Mbp | 32 Mb | 97.7 (3.0) | 30,491 | 5558 | 5386 | 1174 | 94.1 (6.8) |
| **A. saccharum v2** | 571 Mbp | 41 Mb | 96.2 (6.5) | 35,834 | 5366 | 2918 | 1087 | 91.8 (8.5) |
| **A. saccharum v1** | 626 Mbp | 46 Mb | 97.4 (5.6) | 40,074 | 8765 | 6761 | 1190 | 93.1 (7.7) |

\* Complete % (Duplicate %), Embryophyta 10

The *A. saccharum* draft assembly had a total length of 571 Mbp in 1,194 contigs and an N50 of 805 Kb. BUSCO embryophyta genes were 96.0% complete, with 6.6% of those in duplicate.

Similar to *A. negundo*, polishing slightly reduced the total length and N50 in this genome. The BUSCO duplicate score increased by 0.1%. The BUSCO of the scaffolded genome was 96.2% complete, with 6.5% in duplicate, 0.7% fragmented, and 3.1% missing. The assembly had a total length of 571 Mbp in 160 contigs and an N50 of 41 Mb (Table 1, <u>File S1</u>). Scaffolding resulted in 13 chromosomes that represented 96.8% of the assembled length. The genome size dropped from the published reference of 626 Mbp, a more substantial decrease than seen in *A. negundo*, and below the original kmer-based estimate of 636 Mbp. A total of 35,834 genes were identified, and 29,858 were associated with functional information. The resulting annotation BUSCO score was 91.8% complete, containing 8.5% duplicates (Table 1, <u>File S1</u>).

The first versions of the *Acer* genomes were produced exclusively with deep-coverage PacBio reads (PacBio Sequel II, >100X), and resulted in fairly contiguous references that assembled to chromosome-scale with the addition of HiC libraries (McEvoy et al. 2021). The two new accessions, assembled in a hybrid manner and scaffolded with the published PacBio references as described above, were smaller than the original references (Table 1). The difference in size could be structural variation between the different genotypes, but more likely reflects some of the differences in read inputs and methodology (i.e. assembler). *A. negundo* was most similar to its original genome, with an improved duplication rate, and the missing genes identified (∼3 K) were not specific to the new genome. *A. saccharum* had an increase in BUSCO-estimated duplication and 4 K original genes were not observed in the new assembly. Similar to the assembly, differences in final gene number may partially reflect variation in the BRAKER software used for prediction. Whole-genome alignment between the new and original versions revealed that, of the existing assembled sequence, no major discrepancies were present in either species (Fig. 2a). Links between syntenic regions of the original genomes in both species showed similarities in spite of the larger genome size of *A. saccharum* (Fig. 2b).

**Figure 2.**
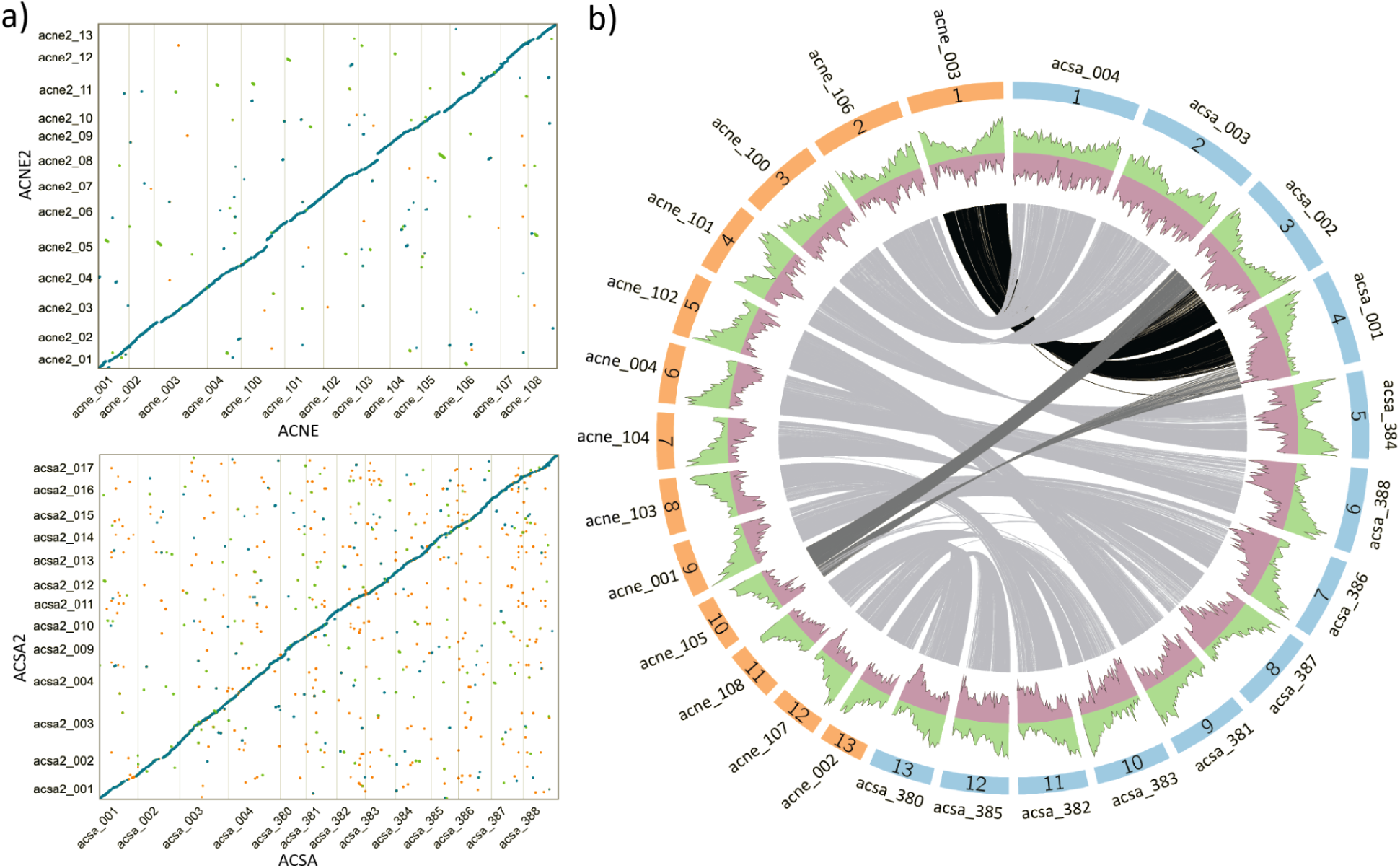
a) Alignment of reference genomes comparing original (x-axis) and new (y-axis) assemblies for *A. negundo* (ACNE) and *A. saccharum* (ACSA). b) Gray lines show blocks of syntenic genes between *A. negundo* (orange) and *A. saccharum* (blue) using the original reference genomes (reprinted from McEvoy et al. (2021)). The green area depicts gene density and purple is repeat frequency.

### Methylation within *Acer*

Long-read sequencing for whole-genome methylation detection and quantification has not yet been widely adopted in plants. *Brassica nigra* was among the first plants on which such analyses were conducted, with ONT reads enabling the demarcation of the centromeres (Perumal et al. 2020); an improved nanopore-sequenced radish genome provided the same centromere-level resolution (Cho et al. 2022). Both centromeres and stress responses were studied in *Gossypium thurberi* and *Gossypium davidsonii* through 5mC and 6mA sites (Yang et al. 2021). The first release of DeepSignal was used with ONT reads to detect novel repeats and characterize the subtelomeres in algae (Chaux-Jukic et al. 2021).

The DeepSignal-Plant tool utilized ONT data from *A. thaliana, Oryza sativa,* and *B. nigra* for development and benchmarking (Ni et al. 2021). This tool represents the first machine-learning approach for plants that can achieve accuracy for all three important 5mC states: CG, CHG, and CHH. The potential roles of methylation in each context is currently being studied in model and non-model systems, but it appears that CG methylation is most often located near and within gene bodies, CHG methylation plays a primary role in silencing TEs, and CHH is responsible for regulating both CG-and CHG-modified TEs (Ni et al. 2021). Studies on transgenerational inheritance of these forms of 5mC modifications in *Arabidopsis* determined that asymmetric CHH must be re-established de novo, while symmetric CG and CHG methylation is maintained (Hsieh, 2016). Using the consensus-based approach from METEORE, we combined DeepSignal-Plant with one of the best implementations of an HMM-based approach, Nanopolish, leveraging the strengths of both to detect methylation across all contexts in ONT sequencing (Yuen et al. 2021).

Independently-assessed methylation levels for CG sites reported by Nanopolish and DeepSignal-Plant were both high relative to the consensus (Fig. 4). DeepSignal-Plant detected the most CG-methylation across all four genomes, as was seen in benchmarking studies comparing DeepSignal (v1) and Nanopolish on *E. coli* and *H. sapiens* data (Ni et al. 2019). The CG levels for both the new and original *A. negundo* and original *A. saccharum* were quite similar (∼70%), and the new *A. saccharum* genome was higher, at 75%. Nanopolish results followed a similar trend, but with slightly lower values of ∼67%, and 71% for the new *A. saccharum* genome. Consensus results for all were around 53%. In particular, the original *A. saccharum* genome was considerably lower, at 34%—close to the CHG value, which was unexpected.

**Figure 3.**
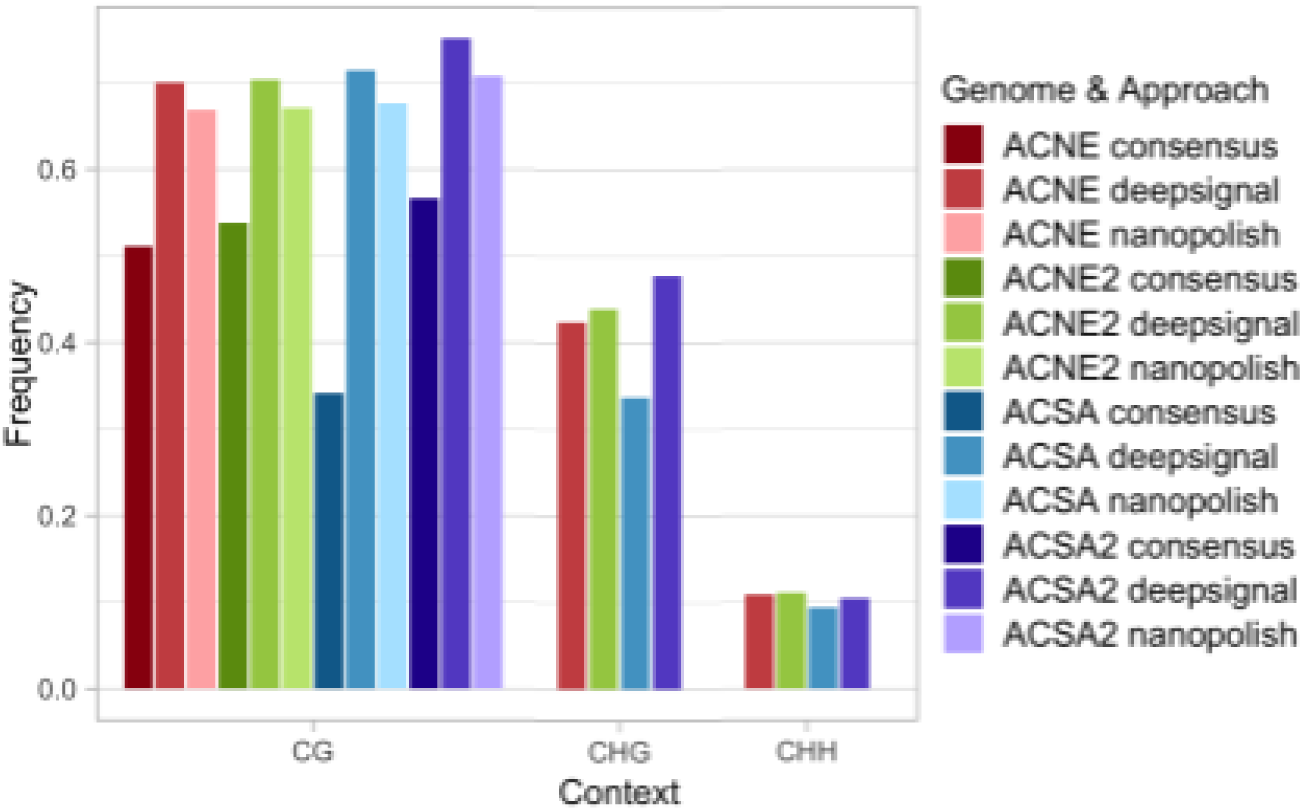
Comparison of global methylation levels by context for DeepSignal-Plant, Nanopolish, and the METEORE-generated random forest consensus.

**Figure 4.**
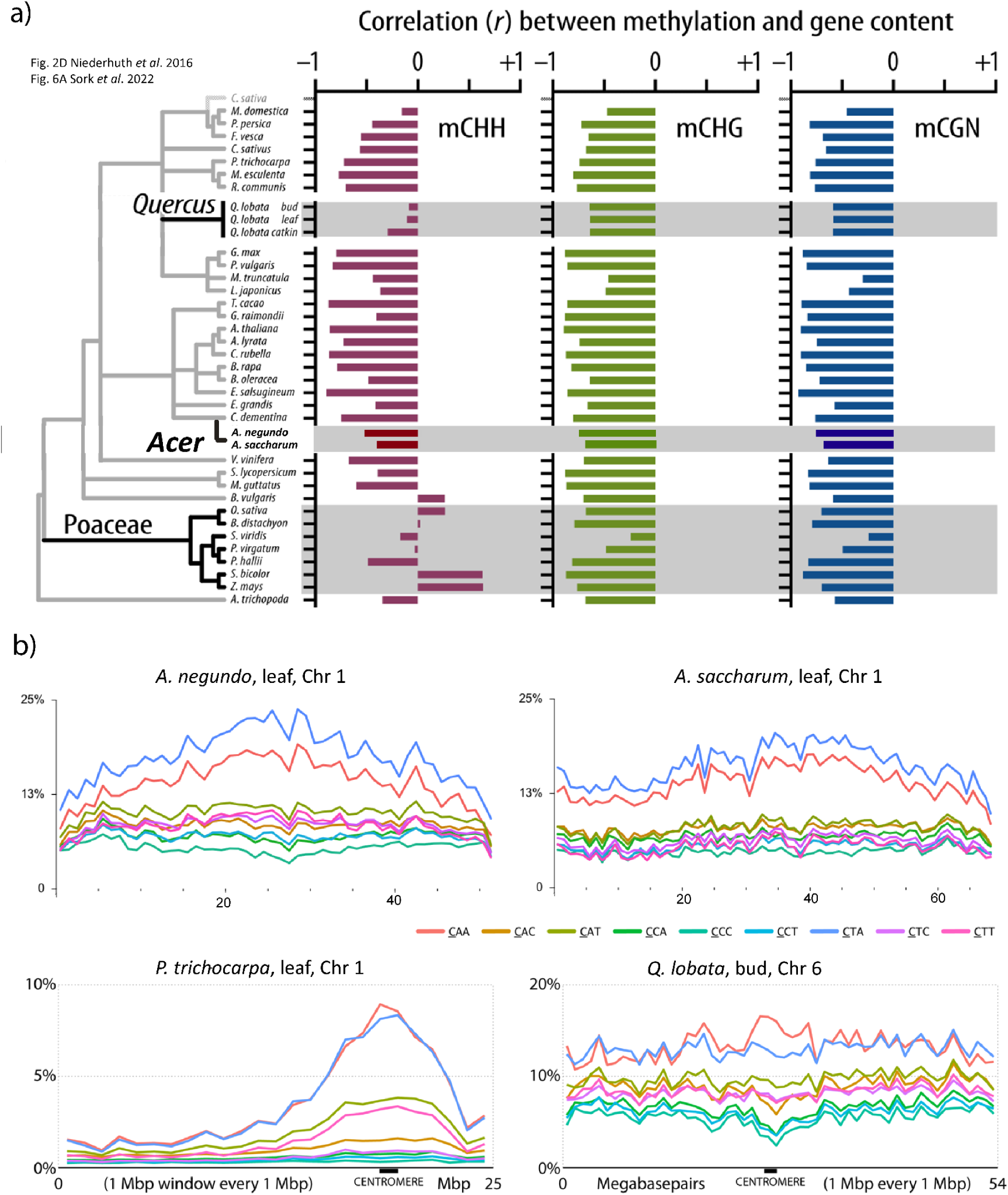
a) Modified from Sork et al. (2022) to add the *Acer* genomes. *Acer* has a negative correlation between methylation and gene content, similar to other species. In contrast, *Quercus* or *Poaceae* are less negatively correlated, or even positively correlated in some species. b) Distribution of CHH methylation by subcontext across the longest chromosomes for *A. negundo* and *A. saccharum*. *Populus* and *Q. lobata*, reprinted from Sork et al. (2022), are shown for comparison of distribution patterns, where *Populus* is localized and *Q. lobata* is generally distributed. Distributions for all chromosomes are in <u>Fig. S1</u>. CHG distributions are in <u>Fig. S2</u> and gene densities plotted with all contexts in 100 Kb windows are in <u>Fig. S3</u>.

The CGH values estimated by DeepSignal-Plant were in the 40% range but, again, the original *A. saccharum* genome value was lower than those of the other genomes, at 33%. CHH values were the lowest, and more consistent across all four, ranging from 9 to 11%.

The original *A. saccharum* genome resulted in much lower estimates of methylation compared to the other three genomes. This genome had the lowest retention of sequences during resquiggling, a preliminary step for DeepSignal-Plant that leverages the reference genome to correct base calling inaccuracies (Oxford Nanopore Technologies 2020b). When resquiggling the reads to the original and new *A. negundo* genomes, 21.2% and 18.0% reads were unsuccessfully processed. For *A. saccharum* these numbers were 54.3% and 31.5%, resulting in 43X coverage for the original genome (File S1). The reduction in retained reads could potentially have reduced coverage to levels affecting methylation calling in DeepSignal-Plant or Nanopolish; however, the primary impact was observed by the consensus statistics generated by METEORE (Fig. 4).

Given the reduced methylation levels for the consensus using the original *A. saccharum*, the methylation datasets based on the new genomes were selected as the best representation. As such, these genomes were used for downstream analyses conducted with the METEORE consensus results. Figures displaying results for both new and original genomes are available in the supplementary material (Fig. S1–5).

### Comparative methylomes

Independent estimates from DeepSignal-Plant and Nanopolish were on the higher end of the spectrum of global methylation levels reported for 34 angiosperms in Niederhuth et al. (2016). Consensus results were closer, but still higher than many angiosperms, and similar in total to *Fragaria vesca*, *Manihot esculenta*, *Vitis vinifera*, *Brachypodium distachyon*, and *Setaria viridis*. WGBS in the form of MethylC-Seq was used in the Niederhuth et al. (2016) study, so increased signal detection could result from the long-reads, which are able to span low-complexity regions. A recent study compared short-read and long-read assemblies of the *Brassica* genome with both WGBS and ONT-based approaches (Nanopolish). The direct CG methylation profiling using the ONT reads was strongly correlated (>93%) with their WGBS data for the same accession. In addition, ONT long-reads identified centromeric repeats and accompanying hypermethylated signal (Perumal et al. 2020).

The distribution of methylation, genes, and TEs is of interest due to the implications for evolutionary strategies involving methylation’s regulation of genes, TEs, and the boundary regions between. Many plant species exhibit a pattern of gene density along the arms of the chromosome and TE density in centromeric regions (Comai et al., 2017). As an exception, Sork et al. (2022) demonstrated that *Q. lobata* has a more uniform distribution of genes and CHH methylation along the chromosome arm, similar to several *Poaceae*. The Pearson correlation (R) of genes to methylation for each sequence context was plotted along with the previous analyses (Niederhuth et al. 2016; Sork et al. 2022) (Fig. 5a). In *Acer*, there is a moderate negative correlation between CHH methylation and genes—almost 50%—similar to about half of angiosperms surveyed, while the remaining angiosperm species have stronger negative correlation (∼75%). CHH was the context with least negative correlation of genes to methylation for *Acer*. Less correlation of methylation with genes means more intersection in TE filled intergenic space, lending support to the hypothesis that in some species, the balance of CHH methylation, TE suppression, TE proximity to genes, and gene body methylation could indicate complex strategies playing a role regulating key genes at TE boundary regions, as observed in grasses (Martin, Seymour, and Gaut 2021) and *Oryza* (Gallo-Franco et al. 2022).

**Figure 5.**
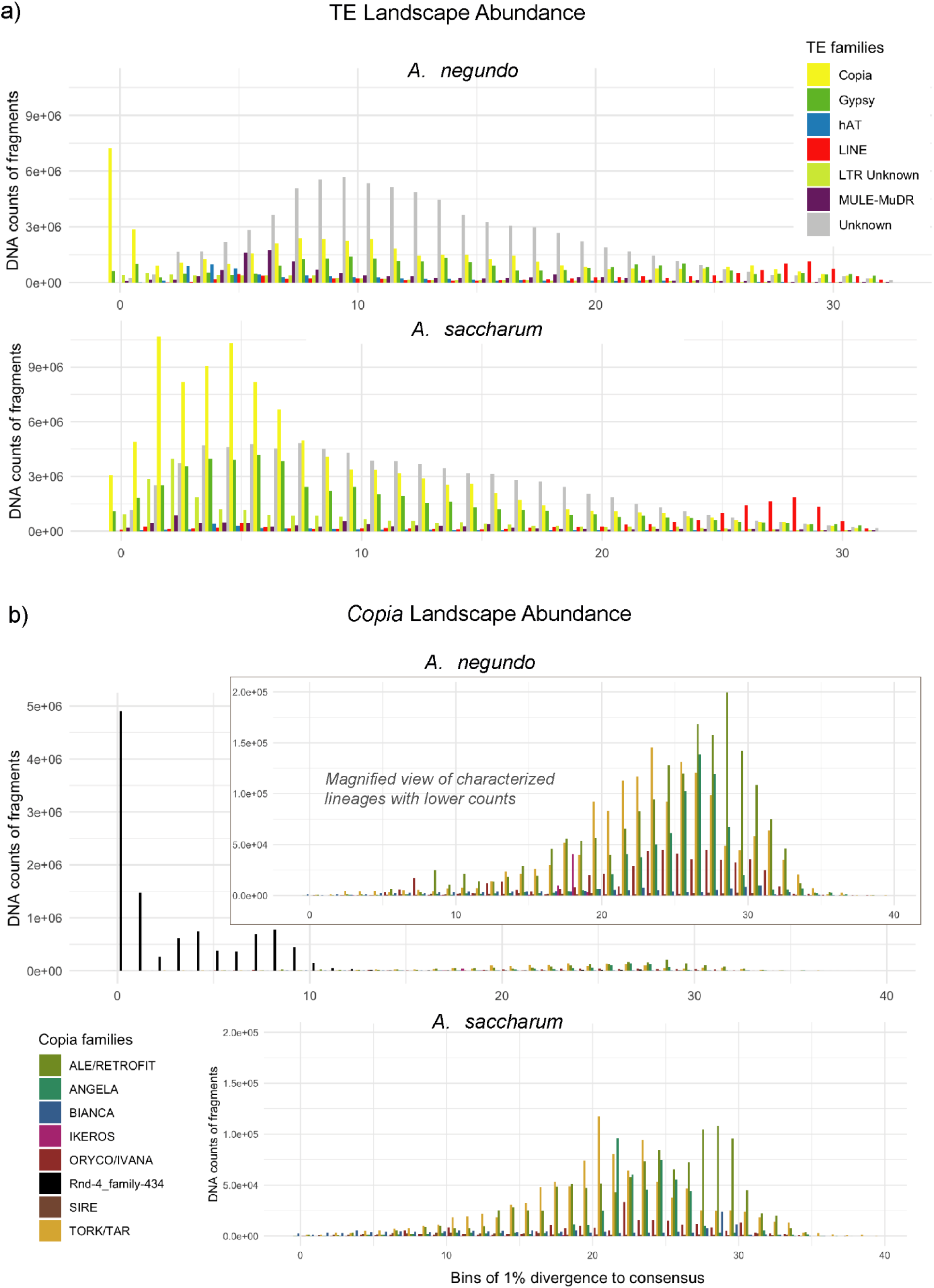
Transposable element abundance for a) predominant repeat families and b) *Copia* subfamilies. The abundance of each repeat class or family is plotted in bins by amount of divergence, with the least divergence representing putative recent activity. In *A. negundo*, a novel *Copia* repeat identified here as *rnd-4_family-434* contributes to the strong recent burst of activity. The other *Copia* subfamilies are provided as an inset for viewability.

Previous analysis of CHH subcontexts across chromosomes detected differences between *Q. lobata*, in which CHH is generally distributed, and *Populus trichocarpa*, in which CHH is localized around the centromere and decreases across the gene-dense chromosome arms (Sork et al. 2022). *Acer* presents an intermediate pattern, particularly *A. saccharum* (Fig. 5b, <u>Fig. S1</u>). *A. saccharum* does not have the same observable enrichment of CHH adjacent to centromeres as *P. trichocarpa*, nor is CHH as generally distributed as in *Q. lobata*. In both *Acer* species, there is a clear pattern for a preference of CTA, followed by CAA, along the lengths of the chromosome arms. Cytosine methyltransferases (CMT) are known to have sequence context preferences. In *Arabidopsis,* CMT2 prefers CHH sites, specifically CAA and CTA. However, little is known about these preferences outside of a handful of model species (Kenchanmane Raju, Ritter, and Niederhuth 2019).

### Repeats

LTR superfamilies *Ty1-copia* and *Ty3-gypsy* typically make up the greatest proportion of TEs in land plants. Insertions of LTRs in and around genes can be responsible for alternative splicing, duplication, recombination, and epigenetic control (Galindo-González et al. 2017). Biotic and abiotic stress, as well as other external stimuli, can lead to TE movement and the rapid proliferation of several subfamilies and may result in polyploidization (Mhiri, Borges, and Grandbastien 2022). The exact mechanisms of activation and repression are not fully understood, but generally involve methylation in any of three sequence contexts. Activation of TEs leads to an initial response of post-transcriptional gene silencing involving siRNA, which is followed by establishment and maintenance of silencing with DNA and histone methylation in RNA-directed DNA methylation (RdDM) pathways. RdDM methylation leads to chromatin modification making the TE inaccessible to transcriptional machinery (Erdmann and Picard 2020). The intersection of methylation and TEs can be informative for understanding control of expression and architectural changes to the genome that may play a role in evolution.

Repeat detection and classification with the LTR database in addition to *de novo* libraries generated by RepeatModeler resulted in a decrease of 9-12% in the previously reported repeat coverages for the original *A. negundo* and *A. saccharum* genomes (McEvoy et al. 2021). The primary reduction was in unknown TEs – those not falling into LTR, DNA, LINE, SINE, or RC classes. LTR coverage decreased slightly (∼0.6%), while the resolution on classification improved to the subfamily level. New and original genomes had small differences in counts and coverage of similar elements, while larger trends remained the same.

The new *A. negundo* genome had 45.2% repeat coverage compared to 45.9% in the original genome. LTRs remained nearly constant at 20% of the genome, with small differences in the family representation. *Copia* had 3,369 unique elements covering 11.2% of the new genome compared to 3,467 covering 10.6% in the original, followed by *Gypsy* with 2,544 unique elements covering 6.4% of the genome and 2,437 covering 7.3% of the original (Table 2, File S1). In addition to *Copia* being the largest superfamily, it also appears to have had a burst of recent activity based on the spike in abundance of copies with low sequence divergence (Fig. 3). The greatest subfamily representation is a novel *Copia* identified here as *rnd-4_family-434*, which covers 0.026% of the genome compared to the next greatest, *ALE/Retrofit* (0.004%), which has more divergent copies.

**Table 2.** Percent of *A. negundo* and *A. saccharum* genomes masked by TE class and *Copia* families.

| TE Class/Family | ACNE | ACSA |
| --- | --- | --- |
| LTR | 19.74% | 31.45% |
| Copia | 11.23% | 18.03% |
| ALE/RETROFIT | 0.004360% | 0.002040% |
| ANGELA | 0.001740% | 0.000849% |
| BIANCA | 0.000261% | 0.000223% |
| IKEROS | 0.000008% | 0.000006% |
| ORYCO/IVANA | 0.00122% | 0.00376% |
| Rnd-4_family-434 | 0.026238% | N/A |
| SIRE | 0.000115% | 0.000101% |
| TORK/TAR | 0.003390% | 0.001490% |
| Gypsy | 6.40% | 9.18% |
| SINE | 0.01% | 0% |
| DNA | 4.69% | 2.58% |
| Caulimovirus | 0.67% | 0.45% |
| hAT | 1.55% | 0.97% |
| MULE-MuDR | 2.69% | 1.35% |
| PIF-Harbinger | 0.19% | 0.16% |
| RC/Helitron | 0.20% | 0.41% |
| LINE | 2.32% | 2.35% |
| Unknown | 18.24% | 14.13% |

*A. saccharum* had greater overall total and LTR repeat coverage, and fewer unique *Copia*/*Gypsy* elements that spanned a greater percentage of the genome compared to *A. negundo*. The new *A. saccharum* was estimated to be 50.9% repetitive while the original genome was estimated at 54.5%. LTRs represented 31% and 37% of the new and original genomes, respectively. There were 3,092 unique elements of *Copia* with 18.0% overall coverage in the new genome compared to 3,015 elements and 22.2% coverage in the original, and only 2,294 unique elements of *Gypsy* with 9.2% coverage in the new genome, and 2,267 elements with 11.0% coverage in the original. *Copia* subfamilies with lower divergence were even more abundant in these genomes meaning there has likely been somewhat recent activity, but the recent burst noted in *A. negundo* was diminished (Fig. 3). *Rnd-4_family-434*, the repeat responsible for the recent burst in *A. negundo*, was not found in *A. saccharum*. In examining LTRs across 69 plant genomes, Orozco-Arias et al (2023) noted that *Copia* lineages (*Oryco*/*Ivana*, *Retrofit*/*Ale*, *Tork*) were frequently represented with high sequence similarity. The *rnd-4_family-434* in *A. negundo* is similar to *Ale/Retrofit* elements and these elements are associated with horizontal transfers (Orozco-Arias et al. 2023).

In most plants, *Gypsy* elements are more abundant and more likely to insert in heterochromatic regions (Vitte et al. 2014), while *Copia* elements are typically more closely associated with genes, more transcriptionally active, and insert in a seemingly random pattern (Galindo-González et al. 2017; Qiu and Ungerer 2018). Among the LTRs presented here, both species were dominated by *Copia* elements, followed by *Gypsy*. Other *Acer* recently characterized, including *A. pseudosieboldianum, A. catalpifolium, A. yangbiense,* and *A. truncatum*, demonstrated the same trend, with greater representation of *Copia* elements (J. Yang et al. 2019; Ma et al. 2020; Yu et al. 2021; X. Li et al. 2022). Exceptions to this include two maples, *A. palmatum* and the polyploid *A. rubrum,* that host more *Gypsy* elements (Lu et al. 2022; Chen et al. 2023). Notable exceptions to the *Gypsy* dominance have also been observed in *Theobroma cacao*, *Vitis vinifera*, *Musa*, *Rhizophora apiculata*, and *Cucumis sativus* (Moisy et al. 2008; Argout et al. 2011; (Wang, Liang, and Tang 2018); Castanera et al. 2019; Pratama, Dwivany, and Nugrahapraja 2021). As more genomes become available, more variation in LTR ratios has been noted within and across genera (Zagorski et al. 2020). It is possible that some of this new variation also reflects improvements in long-read sequencing to resolve and quantify these elements, as seen with *Cucumis melo* (Ruggieri et al., 2018; Castanera et al. 2019). In the *Acers*, despite the fact that the original PacBio reads and the new ONT reads shared similar read N50, the inclusion of the longer ONT reads likely improved the resolution of longer elements. The average length of *Copia* and *Gyspy* elements did in fact increase by several hundred base pairs in *A. negundo,* and while the length of *Copia* elements in *A. saccharum* increased similarly, *Gypsy* elements decreased by 57 bp (File S4).

Methylation across repeat superfamilies was plotted along with results from Sork et al. (2022) for comparison (Fig. 6a, b), but it should be noted that the *Q. lobata* tissue presented was bud, which contains undifferentiated meristem and had three times the CHH methylation compared to catkin and young leaf tissues. Among the LTRs in *Acer*, *Copias* were more frequent than *Gypsy*, with 4.8% more coverage across the *A. negundo* genome and 8.9% in *A. saccharum*. *Copia* elements were slightly less methylated compared to *Gypsy*, particularly in the CG and CHG contexts, with the biggest difference observed in the lower CHG methylation of *Copia* elements in *A. saccharum*. *Copias* were also less methylated in the CHH context of the *Q. lobata* bud tissue (Fig. 6b). In both *Acer*, the reduction of CHH methylation in *Copia* is primarily observed in the up and downstream regions. Interestingly, the greatest frequency of *Copia* CHH methylation for each species is highest in *A. negundo* (∼14%), followed by *A. saccharum* (∼13%), and only ∼7% in *Q. lobata*. Greater methylation is commonly observed in *Gypsy* elements and could be attributed to their tendency to reside in heterochromatic regions ((Wang, Liang, and Tang 2018)).

**Figure 6.**
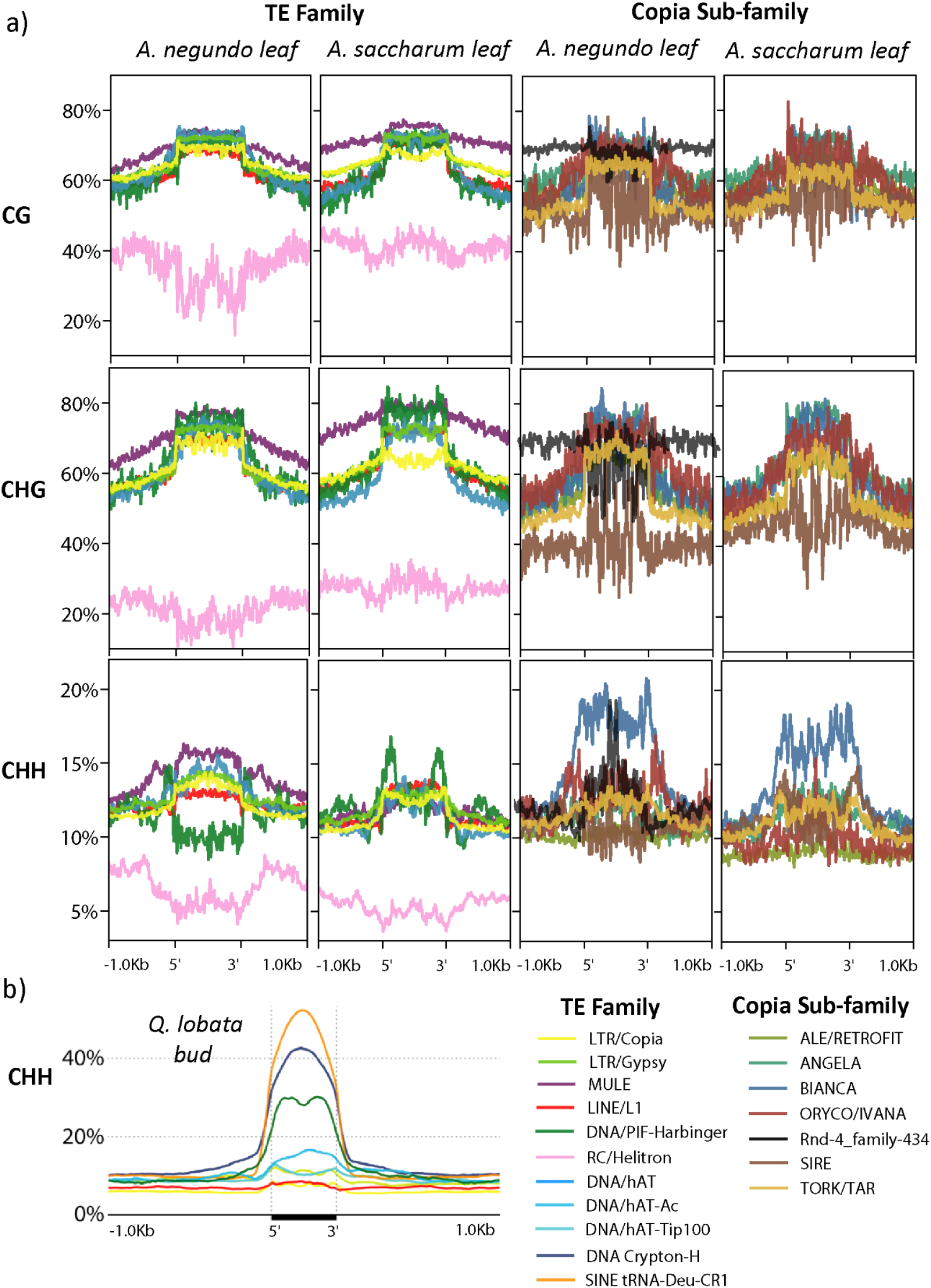
a) Genome-wide methylation distribution by sequence context and TE family. b) Genome-wide CHH distribution (bud tissue) over TE regions, each line representing one of the largest superfamilies of each class. Reprinted from Sork et al. (2022).

Methylation across the characterized and dominant *Copia* subfamilies was also examined (Fig. 6a). Of the shared *Copia* lineages, copy number ranged from *Tork* which, similar to *A. truncatum* (Ma et al. 2020), had the highest representation in *A. saccharum* (5330) and *A. negundo* (5086), to *Sire* which was most abundant in *A. yangbiense* (6624), but was present in far fewer copies in *A. negundo* (770) and *A. saccharum* (1008). In *A. negundo,* a significant contributor to the recent burst of *Copia* activity and the most abundant, *rnd-4_family-434*, had considerably higher CG and CHG methylation in up and downstream regions, and an unusual spike in CHH methylation across the repeat body. In most land plants assessed to date, CHH methylation is observed as a bimodal distribution with most activity in the LTRs themselves (Wang et al. 2018; Noshav et al. 2019; Wang and Baulcombe 2020). CHH methylation among *Bianca*, represented in far fewer copies, was the highest compared to other *Copia* subfamilies in both species and demonstrated the traditional pattern in *A. saccharum* and one less characteristic (more repeat body methylation) in *A. negundo*. In both *Acer*, *Sire* had less methylation than other *Copia* subfamilies in all sequence contexts except *A. saccharum* CHH. In *A. saccharum*, the CHH methylation frequencies of *Oryco/Ivana* and *Ale/Retrofit* were even lower than *Sire*, and while the copy number of *Oryco/Ivana* was also lower, *Ale/Retrofit* was interesting in its overall prevalence in the genome, with 4035 and and 3488 divergent copies for *A. negundo* and *A. saccharum*, respectively. This pattern of high *Ale* sequence diversity and more even distribution through the genome was observed in a population level assessment of *Setaria italica* (Suguiyama et al. 2019). The low CHH methylation observed for Ale/Retrofit elements could be a reflection of sufficient suppression by CG and CHG elements, or it could be ascribed to the inherent structure of these TEs which may influence methylation levels. In *Brachypodium distachyon*, GC content was low in *Sire* and *Bianca*, which was reflected in lower CG and CHG levels, but high in *Ale/Retrofit* with GC content that was decreasing with age due to deamination of cytosines (Stritt et al. 2020).

Studies on CHH methylation in *Arabidopsis* have characterized the unique pathways responsible, and their TE targets (Bouyer et al., 2017). The CMT2 pathway targets the LTRs (primarily *Gypsy* and *Copia*) (Sasaki et al. 2019) and the RdDM pathway targets Class I TEs, specifically RC/*Helitron* and *MULE-MuDR*. In *Acer*, CHH methylation in DNA transposons was strongly increased (Fig. 6a), pointing to a functional role of asymmetric methylation in DNA transposon silencing (Zakrzewski et al. 2017). Among DNA transposons, DNA/*MULE-MuDR* elements were most highly methylated, except for the CHH context in *A. saccharum*, where it was similar in frequency to other TE families. *MULE-MuDR* elements contribute to genome crossovers as they are prone to meiotic double-strand breaks (Underwood and Choi 2019). The insertion of MuDRs in maize increases meiotic crossover rates and these primarily occur in regions of open chromatin (Liu et al. 2009). The loss of CG and non-CG methylation over MuDr elements in *Arabidopsis* mutants led to an increase in crossovers that were enriched in proximity to immunity genes of large and diverse families with extensive variation (Choi et al. 2018). DNA/*PiF-Harbinger* elements were also highly methylated across the gene body with the exception of CHH methylation in *A. negundo*. CHH methylation of *PiF-Harbinger* has a bimodal distribution, with peaks flanking the element in both *Acer*s, similar to *Q. lobata* (Fig. 6b), but the overall frequency of CHH methylation is much higher in oak (∼30%) which may be a tissue specific pattern. *PiF-Harbinger* elements only cover a very small portion of the *Acer* genomes at ∼0.18%, but they are widespread throughout plant lineages, at times in great abundance (∼1%) relative to most DNA transposons as observed in *Arabidopsis thaliana*, *Oryza sativa*, and *Brassica oleracea* (Zhang et al. 2004). They have been associated with important adaptive roles, including intronic *Harbinger* elements that contain MYB elements that regulate stress response to light in *Solanum lycopersicum* (Deneweth, Van de Peer, and Vermeirssen 2022), and a *Harbinger*-derived gene that coordinates histone modifications that alter panicle number and grain size in *O. sativa* (Mao et al. 2022).

Among non-LTR retroelements, *SINE*s were heavily methylated (>40% for CHH) in *Quercus* bud tissue (Fig. 6b) but not frequently identified in the *Acer* for comparison (File S1). *SINEs* are derived from tRNA and demonstrate low sequence similarity which may be responsible for their poor annotation (Wenke et al. 2011). Recent analysis across eight grass genomes examined patterns in mCHH islands and found significant enrichment of these islands in DNA transposons (DNA/*MULE-MuDR, PiF-Harbinger)* as well as non-LTR retroelements (SINEs) (Martin, Seymour, and Gaut 2021).

Methylation frequencies across gene regions in *Acer* were somewhat different than that observed in *Q. lobata*, but there is a spectrum of variation observed across angiosperms, as seen in Niederhuth et al. (2016) (Fig. 4a, Fig. 7a, b). The two *Acer* species were very similar, but *A. negundo*, which has the smaller genome, had a slightly higher frequency of CG. The higher frequencies across genes in the original genomes compared to the new genomes may be due to differences in average gene length, as seen in *Q. lobata* (Fig. 7a, <u>Fig. S4</u>). Both *Acer* species also shared the same sharp increase in CHH just upstream of the translation start site, which was greater in *A. negundo*.

**Figure 7.**
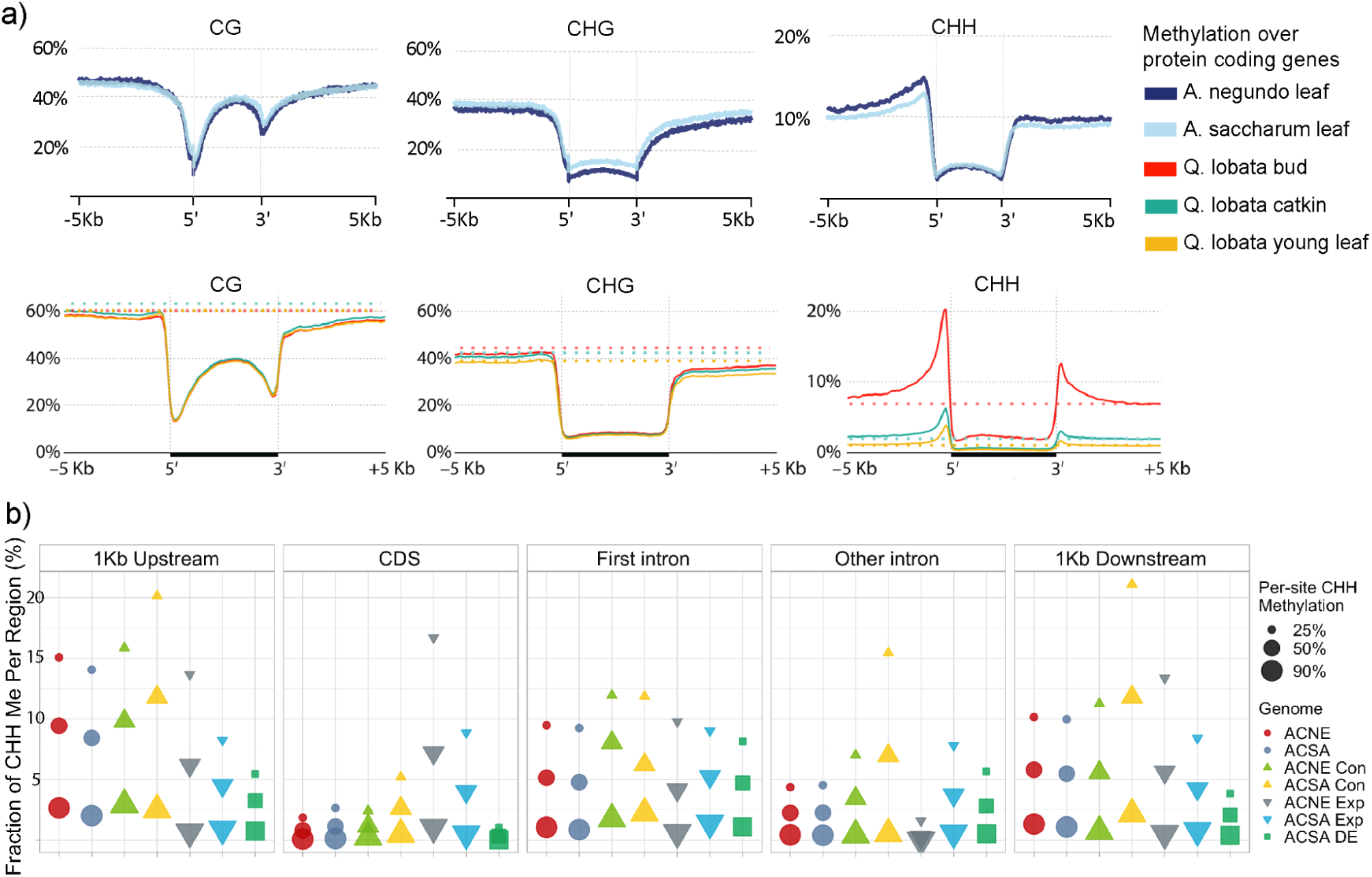
a) Methylation frequency distribution across protein coding regions, shown by genome, tissue type, and sequence context, b) Methylation frequencies at different levels (25%, 50%, 90%), in different genic regions (1 Kb upstream, CDS, first intron, other introns, and, 1 Kb downstream), for different genome-wide subsets. Abbreviations are, ACNE/ACSA: genome-wide means, Con: rapidly contracting genes, Exp: rapidly expanding genes, DE: differentially expressed genes. Exp and Con genes are from McEvoy et al. (2021), as well as DE, where genes were differentially expressed in response to calcium and aluminum treatment. Of the original 245 DE genes identified from stem tissue samples, 240 are presented here with methylation calculated from leaf tissue.

Focusing on CHH levels across genic regions, higher frequencies were found in portions of the upstream flanking region (Fig. 7a, 7b). The next largest fraction was in the first intron, where both new *Acer* genomes had a significant drop in the number of methylated sites, especially for those at 90% methylation (Fig. 7b). This pattern was seen in other introns, but at lower levels, while the levels in downstream flanking regions were consistent across frequencies. In certain plant genes—often highly expressed genes—the first intron contains regulatory elements, though it is observed much less frequently than upstream promoters (Rose 2018).

### Methylome and gene regulation

By combining distributions of methylation, select TEs, and gene density, trends amongst the elements can be observed. In Fig. 8, each row contains chromosomes that are largely syntenic between *A. negundo* (ACNE) and *A. saccharum* (ACSA), as seen in Fig. 2b. Regions of low gene density are co-located with peaks of LTRs and methylation in all contexts, including CHH. It is likely these contain centromeres and pericentromeric regions. Included in the gene density track (Fig. 8) are genes from families previously identified as significantly expanding (26 in *A. negundo*, 99 in *A. saccharum*) or significantly contracting (52 in *A. negundo*, 18 in *A. saccharum*) when evaluated in terms of gene family evolution across 22 land plant species (McEvoy et al. 2021). Genes from rapidly evolving families are found in every chromosome except acsa2_011, though some contain many more than others (<u>Fig. S5</u>).

**Figure 8.**
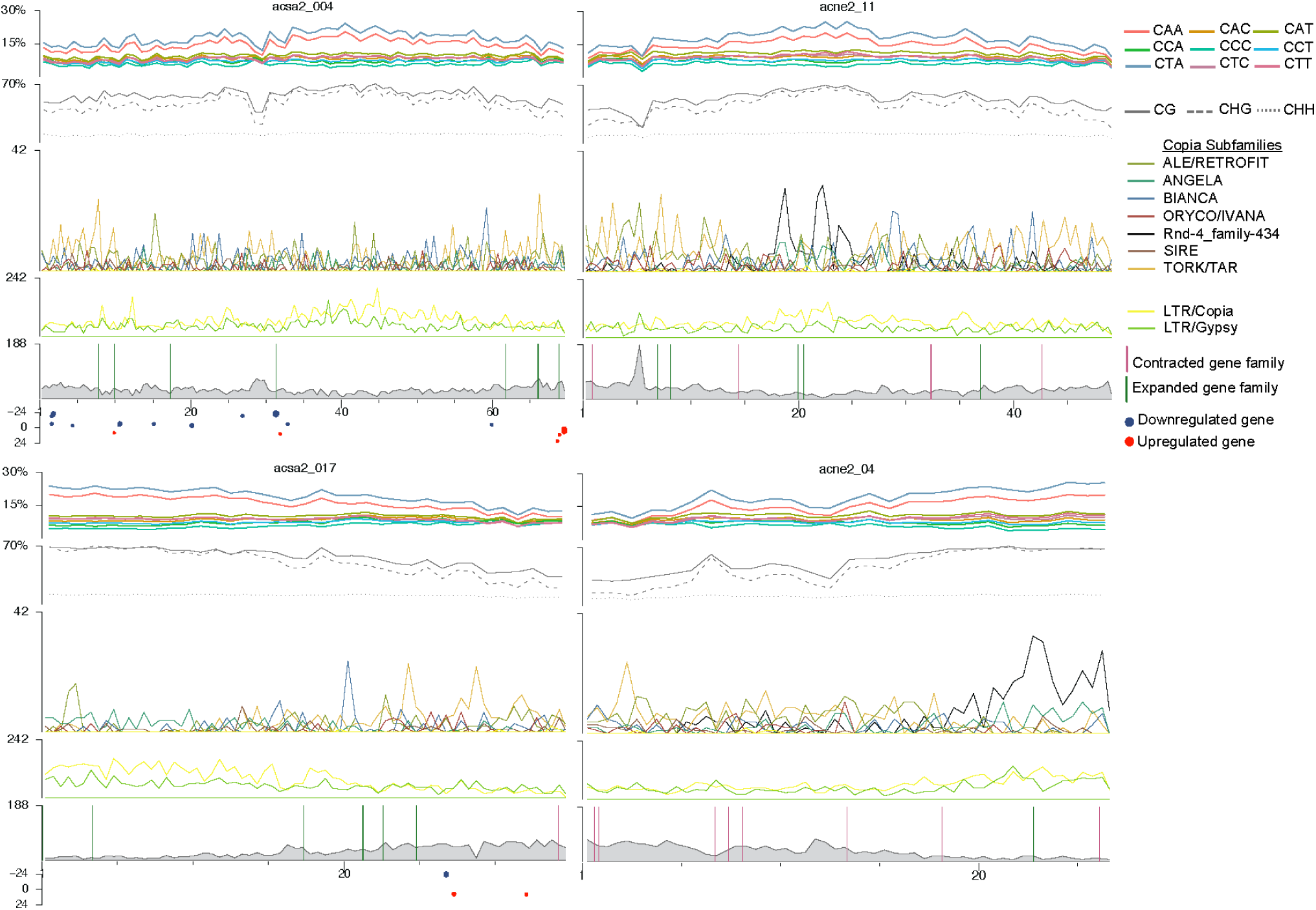
Distribution of gene density, repeat families, methylation by context, CHH methylation subcontexts, gene family dynamics, and gene expression results for select chromosomes. Middle row, right, shows multiple gene family expansions (lime green vertical bars) in *A. saccharum*. Bottom row shows gene family expansions and contraction (purple bars) in *A. negundo*. Gene expression results are from aluminum and calcium treatments at Hubbard Brook Experimental Forest as detailed in McEvoy et al. (2021). X-axis indicates log2 fold change, while dot size represents the *p*-adjusted value.

The particular subset of chromosomes shown in Fig. 8 provides a few different examples of distribution patterns and rapid gene family evolution in chromosomes. The top set of syntenic chromosomes, acsa2_004 (*A. saccharum*) and acne2_11 (*A. negundo*), shows a generally even distribution of gene density rather than a concentration of genes in the chromosome arms. The *A. negundo* chromosome (acne2_11) had the most genes from expanded families, and it also contains some genes from contracted families. There is a notable drop in methylation over a narrow peak of gene density in both species that is greater in *A. negundo*. *A. negundo* also has spikes of high frequency *rnd-4_family-434* that correspond to the greatest methylation frequencies and lowest gene densities. *Rnd-4_family-434*, the Copia subfamily with the burst of recent activity, does not appear to be generally associated with rapidly evolving genes (<u>Fig. S5</u>), but there are instances where the two overlap, as shown in acne2_11, where two genes from rapidly expanding families fall between the two spikes of *rnd-4_family-434*.

The second set of chromosomes, acsa2_017 and acne2_04, are also syntenic, but inverted, and gene density is concentrated in one arm of each species. In this distribution, *rnd-4_family-434* is concentrated at the opposite end of the *A. negundo* chromosome where gene density is the lowest and methylation the greatest. This distribution pattern, weighted towards one chromosome arm, tends to be more common genome-wide in both species (<u>Fig.</u> <u>S5</u>). The *A. negundo* chromosome has several genes from contracted families and one from expanded, while *A. saccharum* is the inverse, which fits the predominant direction of rapidly evolving gene families for each species. In the previous *Acer* study, *A. negundo* was shown to have more families associated with rapid contraction, and here, expanded gene families show an increase in CHH methylation in the gene body (coding regions) in both species, but it is greater in *A. negundo*. *A. saccharum* primarily had expanding families, and here, genes from contracted families have distinct increases in CHH methylation upstream, downstream, and in the intronic space, regardless of position, particularly in *A. saccharum* (Fig. 7b).

Several studies have examined the relationship between methylation changes, metal toxicity, and the associated nutrient stress, in both model and non-model plant systems. One such study examined the aquatic plant *Hydrilla verticillata* and noted demethylation in response to copper, as well as blocked ROS interactions that could cause remethylation (Shi et al. 2017). In *Hibiscus cannabinus*, increased methylation was associated with increasing chromium levels (Ding et al. 2016). In *Arabidopsis halleri*, cadmium treatments increased CG methylation of genes responsible for symmetric methylation (Galati et al. 2021). In *O. sativa*, a model for nutrient stress, the expression of Heavy Metal Transporting ATPases (HMAs) was modified and retained over generations and modulated by the methylation of specific TEs (Cong et al. 2019). Gallo-Franco et al. (2022) found that aluminum tolerant and susceptible varieties of *O. sativa* and *Oryza glumaepatula* each had unique differentially methylated regions, and each species was associated with different TEs potentially affecting regulation of these regions. While relationships between environmental stress, methylation, and increased proximal TE activity have been observed in numerous studies, factors can be complex and potentially involve selection of variants affecting methylation mechanisms or chromatin features, such as the *Copia ONSEN* in both *Arabidopsis* and *Oryza*. This TE inserts near histone variant H2A.Z, the deposition of which can be regulated by environment (Baduel and Quadrana 2021). While our study does not lend itself to proper examination of species-specific patterns of methylation that can be correlated to gene expression changes, the potential for gene regulation among candidate genes previously identified in RNA-Seq experiments in *A. saccharum* could be examined.

Gene expression results focused on candidates associated with nutrient stress (or heavy metal toxicity) were mapped to the new *A. saccharum* genome (240 in total; Fig. 8). A total of 115 genes were downregulated in trees grown in aluminum-amended plots while 130 genes were up-regulated (in both cases, compared with trees grown in calcium-amended or control plots). These genes are distributed across all chromosomes at different densities. To further examine the patterns of methylation in all three contexts, the 240 differentially expressed genes were compared with the patterns observed for the mean of all genes in Fig. 7b and File S4. The subset of differentially expressed genes appeared enriched for CG and CHG methylation in the upstream regions when compared with the full gene space. The most noticeable difference, however, was the higher peak of CHH methylation frequency immediately preceding the translation start site in the promoter region of differentially expressed genes (<u>Fig. S4</u>, Fig. 7b) which has been observed in grasses and other angiosperms (Martin, Seymour, and Gaut 2021). Values here peak at around 16% compared to 12.5% for the whole genome mean. This value is comparable to the CHH methylation observed in *Q. lobata* bud tissue relative to catkins or young leaves, which have a greater portion of meristematic tissue, as mentioned in Sork et al. (2022), and are presumably undergoing developmental processes. Increased methylation could be a sign of gene networks requiring more flexibility in expression to meet the challenges of development or biotic and abiotic stress (Lang et al., 2017).

## CONCLUSION

Plants are dependent on methylation to regulate both short and long-term adaptive strategies. Best practices to detect this methylation leading to an understanding of integrated epigenomic data are important for a full understanding of the genomic mechanics in effect. To extend previous work on DNA methylation detection and integration, this study developed two improved reference genomes for two new accessions of *A. saccharum* and *A. negundo*. These hybrid assemblies benefitted from the inclusion of deep ONT coverage to both resolve long repeat elements and detect methylation. Methylation calling was conducted with ONT sequencing data and a custom pipeline. This pipeline, which leverages super accuracy base calling and detection of false positives, has broader application for other non-model plant genomes. Methylation frequencies and distributions were compared with other recent plant methylomes and revealed differences among the species where *A. saccharum* and *A. negundo* have a lower correlation of methylation and gene content than many angiosperms and fall in the middle of the spectrum of gene dense arms with low methylation of *P. trichocarpa* and the generalized distribution of methylation in *Q. lobata* and *Poaceae*. In both *Acer* species, the repeat landscape was dominated by LTR/*Copia* elements. A recent burst of a novel *Copia* in *A. negundo* showed higher than average CG and CHG methylation up and downstream and an unusual pattern of strong CHH methylation in the repeat body. Gene family dynamics integrated with methylation data noted that individual genes in contracted families were associated with strong CHH methylation upstream, downstream, and in intronic regions. Expanded gene families had more CHH methylation in coding regions than contracted families. Among candidate genes previously associated with nutrient stress, patterns of methylation were variable, but had increased upstream methylation observed in all three contexts. Further investigations require parallel expression and tissue-specific studies, as well as pan-genome (population scale) analysis to understand how the methylome is contributing to genome evolution.

## DATA AVAILABILITY

New and original *Acer saccharum* sequencing, assemblies, and annotations are available in BioProject PRJNA748028. *Acer negundo* data is available in BioProject PRJNA750066. Scripts are available in the Plant Computational Genomics GitLab, AcerMethylation repository https://gitlab.com/PlantGenomicsLab/acermethylation/

## ACKNOWLEDGEMENTS

We would like to thank the Arnold Arboretum of Harvard University for providing leaf samples that were difficult to acquire during the pandemic. We thank the Institute for Systems Genomics Center for Genome Innovation at the University of Connecticut for molecular resources and support. We would also like to thank the Computational Biology Core for access to HPC resources. We would also like to thank members of the University of Connecticut MCB and EEB departments, including Christine McCann who advised on extractions and library prep, and the many who provided collaborative brainstorming on methylation calling methods, including Dr. Savannah Hoyt, Gabby Hartley, Michelle Neitzey, Vidya Vuruputoor, and Dr. Ross Whetten at NCSU. We would also like to thank Dr. Pamela Diggle, Dr. Louise Lewis, Dr. Fay-Wei Li, Dr. Elizabeth Jockusch, and Dr. Yaowu Yuan for their comments on the draft manuscript.

## SUPPLEMENTARY MATERIAL

### Supplementary Files

File S1: Assembly statistics, annotation statistics, summaries of repeat coverage by class and family, tombo resquiggle summaries

https://gitlab.com/PlantGenomicsLab/acermethylation/-/blob/main/manuscript/supplemental/FileS1-summaries.xlsx

### Supplementary Figures

Figure S1: Distribution of CHH methylation by subcontext across all chromosomes (1 Mbp window every 1 Mbp) in new and original *A. negundo* (acne) and *A. saccharum* (acsa) genomes.

Figure S2: Distribution of CHG methylation by subcontext across all chromosomes (1 Mbp window every 1 Mbp) in new and original *A. negundo* (acne) and *A. saccharum* (acsa) genomes.

Figure S3: Gene densities (red) with CG (blue), CHG (green), and CHH (maroon) contexts in 100 Kb windows across new and original *A. negundo* (acne) and *A. saccharum* (acsa) genomes.

Figure S4: a) Methylation frequency distribution across protein coding regions, 5’ to 3’, shown by assembly and sequence context for new *A. negundo* (acne2) and *A. saccharum* (acsa2) genomes. Top row includes results from whole-genome mean, second row is genes from rapidly expanding gene families, third row is genes from rapidly contracting families. Bottom row shows methylation frequency distribution across 240 (of the original 245) genes differentially expressed in response to calcium and aluminum treatments in stem as seen in McEvoy et al. (2021).

Figure S5: Chromosome-level distribution of gene density, LTR repeat families and Copia subfamilies, methylation by context, CHH methylation subcontexts, gene family dynamics, and gene expression results for select chromosomes. New *A. negundo* (acne2) and *A. saccharum* (acsa2) are shown. Genes from expanded gene families are shown as lime green vertical bars, and those from contracted, as purple bars. *A. saccharum* gene expression results (red and blue dots) are from aluminum and calcium treatments at Hubbard Brook Experimental Forest as detailed in McEvoy et al. (2021). X-axis indicates log2 fold change, while dot size represents the p-adjusted value.

## REFERENCES

1. Alonge, Michael, Ludivine Lebeigle, Melanie Kirsche, Sergey Aganezov, Xingang Wang, Zachary B. Lippman, Michael C. Schatz, and Sebastian Soyk. 2021. “Automated Assembly Scaffolding Elevates a New Tomato System for High-Throughput Genome Editing.” bioRxiv. https://doi.org/10.1101/2021.11.18.469135.

2. Argout, Xavier, Jerome Salse, Jean-Marc Aury, Mark J. Guiltinan, Gaetan Droc, Jerome Gouzy, Mathilde Allegre, et al. 2011. “The Genome of Theobroma Cacao.” Nature Genetics 43 (2): 101–8.

3. Baduel, P., & Quadrana, L. (2021). Jumpstarting evolution: How transposition can facilitate adaptation to rapid environmental changes. Current Opinion in Plant Biology, 61, 102043.

4. Bishop, Daniel A., Colin M. Beier, Neil Pederson, Gregory B. Lawrence, John C. Stella, and Timothy J. Sullivan. 2015. “Regional Growth Decline of Sugar Maple (Acer Saccharum) and Its Potential Causes.” Ecosphere 6 (10): art179.

5. Brůna, Tomáš, Katharina J. Hoff, Alexandre Lomsadze, Mario Stanke, and Mark Borodovsky. 2021. “BRAKER2: Automatic Eukaryotic Genome Annotation with GeneMark-EP+ and AUGUSTUS Supported by a Protein Database.” NAR Genomics and Bioinformatics 3 (1): lqaa108.

6. Bouyer, D., Kramdi, A., Kassam, M., Heese, M., Schnittger, A., Roudier, F., & Colot, V. (2017). DNA methylation dynamics during early plant life. Genome Biology, 18(1), 179.

7. Caballero, Madison, and Jill Wegrzyn. 2019. “gFACs: Gene Filtering, Analysis, and Conversion to Unify Genome Annotations Across Alignment and Gene Prediction Frameworks.” Genomics, Proteomics & Bioinformatics 17 (3): 305–10.

8. Castanera, Raúl, Valentino Ruggieri, Marta Pujol, Jordi Garcia-Mas, and Josep M. Casacuberta. 2019. “An Improved Melon Reference Genome with Single-Molecule Sequencing Uncovers a Recent Burst of Transposable Elements With Potential Impact on Genes.” Frontiers in Plant Science 10: 1815.

9. Cavrak, Vladimir V., Nicole Lettner, Suraj Jamge, Agata Kosarewicz, Laura Maria Bayer, and Ortrun Mittelsten Scheid. 2014. “How a Retrotransposon Exploits the Plant’s Heat Stress Response for Its Activation.” PLoS Genetics 10 (1): e1004115.

10. Chaux-Jukic, Frédéric, Samuel O’Donnell, Rory J. Craig, Stephan Eberhard, Olivier Vallon, and Zhou Xu. 2021. “Architecture and Evolution of Subtelomeres in the Unicellular Green Alga Chlamydomonas Reinhardtii.” Nucleic Acids Research 49 (13): 7571–87.

11. Chen, Shifu, Yanqing Zhou, Yaru Chen, and Jia Gu. 2018. “Fastp: An Ultra-Fast All-in-One FASTQ Preprocessor.” Bioinformatics 34 (17): i884–90.

12. Chen, Z., Lu, X., Zhu, L., Afzal, S. F., Zhou, J., Ma, Q., Li, Q., Chen, J., & Ren, J. (2023). Chromosomal-level genome and multi-omics dataset provides new insights into leaf pigmentation in Acer palmatum. International Journal of Biological Macromolecules, 227, 93–104.

13. Cho, Ara, Hoyeol Jang, Seunghoon Baek, Moon-Jin Kim, Bomi Yim, Sunmi Huh, Song-Hwa Kwon, Hee-Ju Yu, and Jeong-Hwan Mun. 2022. “An Improved Raphanus Sativus Cv. WK10039 Genome Localizes Centromeres, Uncovers Variation of DNA Methylation and Resolves Arrangement of the Ancestral Brassica Genome Blocks in Radish Chromosomes.” Theoretical and Applied Genetics 135 (5): 1731–50.

14. Choi, K., Zhao, X., Tock, A. J., Lambing, C., Underwood, C. J., Hardcastle, T. J., Serra, H., Kim, J., Cho, H. S., Kim, J., Ziolkowski, P. A., Yelina, N. E., Hwang, I., Martienssen, R. A., & Henderson, I. R. (2018). Nucleosomes and DNA methylation shape meiotic DSB frequency in Arabidopsis thaliana transposons and gene regulatory regions. Genome Research, 28(4), 532–546.

15. Ci, Dong, Yuepeng Song, Qingzhang Du, Min Tian, Shuo Han, and Deqiang Zhang. 2016. “Variation in Genomic Methylation in Natural Populations of Populus Simonii Is Associated with Leaf Shape and Photosynthetic Traits.” Journal of Experimental Botany 67 (3): 723–37.

16. Comai, L., Maheshwari, S., & Marimuthu, M. P. A. (2017). Plant centromeres. Current Opinion in Plant Biology, 36, 158–167.

17. Cong, Weixuan, Yiling Miao, Lei Xu, Yunhong Zhang, Chunlei Yuan, Junmeng Wang, Tingting Zhuang, et al. 2019. “Transgenerational Memory of Gene Expression Changes Induced by Heavy Metal Stress in Rice (Oryza Sativa L.).” BMC Plant Biology 19 (1): 282.

18. Deneweth, J., Van de Peer, Y., & Vermeirssen, V. (2022). Nearby transposable elements impact plant stress gene regulatory networks: a meta-analysis in A. thaliana and S. lycopersicum. BMC Genomics, 23(1), 18.

19. Ding, Han, Guodong Wang, Lili Lou, and Jinyin Lv. 2016. “Physiological Responses and Tolerance of Kenaf (Hibiscus Cannabinus L.) Exposed to Chromium.” Ecotoxicology and Environmental Safety 133 (November): 509–18.

20. Erdmann, Robert M., and Colette L. Picard. 2020. “RNA-Directed DNA Methylation.” PLoS Genetics 16 (10): e1009034.

21. Fan, Xiaoru, Lirun Peng, and Yong Zhang. 2022. “Plant DNA Methylation Responds to Nutrient Stress.” Genes 13 (6): 992.

22. Flynn, Jullien M., Robert Hubley, Clément Goubert, Jeb Rosen, Andrew G. Clark, Cédric Feschotte, and Arian F. Smit. 2020. “RepeatModeler2 for Automated Genomic Discovery of Transposable Element Families.” Proceedings of the National Academy of Sciences of the United States of America 117 (17): 9451–57.

23. Galati, Serena, Mariolina Gullì, Gianluigi Giannelli, Antonella Furini, Giovanni DalCorso, Rosaria Fragni, Annamaria Buschini, and Giovanna Visioli. 2021. “Heavy Metals Modulate DNA Compaction and Methylation at CpG Sites in the Metal Hyperaccumulator Arabidopsis Halleri.” Environmental and Molecular Mutagenesis 62 (2): 133–42.

24. Galindo-González, Leonardo, Corinne Mhiri, Michael K. Deyholos, and Marie-Angèle Grandbastien. 2017. “LTR-Retrotransposons in Plants: Engines of Evolution.” Gene 626 (August): 14–25.

25. Gallo-Franco, J. J., Ghneim-Herrera, T., Tobar-Tosse, F., Romero, M., Chaura, J., & Quimbaya, M. (2022). Whole-genome DNA methylation patterns of Oryza sativa (L.) and Oryza glumaepatula (Steud) genotypes associated with aluminum response. Plant Direct, 6(8), e430.

26. Gel, Bernat, and Eduard Serra. 2017. “karyoploteR: An R/Bioconductor Package to Plot Customizable Genomes Displaying Arbitrary Data.” Bioinformatics 33 (19): 3088–90.

27. Gouil, Quentin, and Andrew Keniry. 2019. “Latest Techniques to Study DNA Methylation.” Essays in Biochemistry 63 (6): 639–48.

28. Hart, Alexander J., Samuel Ginzburg, Muyang (sam) Xu, Cera R. Fisher, Nasim Rahmatpour, Jeffry B. Mitton, Robin Paul, and Jill L. Wegrzyn. 2018. “EnTAP: Bringing Faster and Smarter Functional Annotation to Non-Model Eukaryotic Transcriptomes.” bioRxiv, April, 3078

29. He, Li, Huan Huang, Mariem Bradai, Cheng Zhao, Yin You, Jun Ma, Lun Zhao, Rosa Lozano-Durán, and Jian-Kang Zhu. 2022. “DNA Methylation-Free Arabidopsis Reveals Crucial Roles of DNA Methylation in Regulating Gene Expression and Development.” Nature Communications 13 (1): 1335.

30. He, Li, Wenwu Wu, Gaurav Zinta, Lan Yang, Dong Wang, Renyi Liu, Huiming Zhang, et al. 2018. “A Naturally Occurring Epiallele Associates with Leaf Senescence and Local Climate Adaptation in Arabidopsis Accessions.” Nature Communications 9 (1): 460.

31. Hofmeister, Brigitte T., Johanna Denkena, Maria Colomé-Tatché, Yadollah Shahryary, Rashmi Hazarika, Jane Grimwood, Sujan Mamidi, et al. 2020. “A Genome Assembly and the Somatic Genetic and Epigenetic Mutation Rate in a Wild Long-Lived Perennial Populus Trichocarpa.” Genome Biology 21 (1): 259.

32. Hsieh, P.-H. (2016). *Maintenance and Inheritance of DNA Methylation in Arabidopsis* [UC Berkeley]. https://escholarship.org/uc/item/9r85v5k7

33. Jullien, Pauline E., Daichi Susaki, Ramesh Yelagandula, Tetsuya Higashiyama, and Frédéric Berger. 2012. “DNA Methylation Dynamics during Sexual Reproduction in Arabidopsis Thaliana.” Current Biology: CB 22 (19): 1825–30.

34. Kenchanmane Raju, Sunil K., Eleanore Jeanne Ritter, and Chad E. Niederhuth. 2019. “Establishment, Maintenance, and Biological Roles of Non-CG Methylation in Plants.” Essays in Biochemistry 63 (6): 743–55.

35. Kim, Daehwan, Li Song, Florian P. Breitwieser, and Steven L. Salzberg. 2016. “Centrifuge: Rapid and Sensitive Classification of Metagenomic Sequences.” Genome Research 26 (12): 1721–29.

36. Kumar, Suresh, and Trilochan Mohapatra. 2021. “Dynamics of DNA Methylation and Its Functions in Plant Growth and Development.” Frontiers in Plant Science 12 (May): 596236.

37. Lang, Z., Wang, Y., Tang, K., Tang, D., Datsenka, T., Cheng, J., Zhang, Y., Handa, A. K., & Zhu, J.-K. (2017). Critical roles of DNA demethylation in the activation of ripening-induced genes and inhibition of ripening-repressed genes in tomato fruit. Proceedings of the National Academy of Sciences of the United States of America, 114(22), E4511–E4519.

38. Langmead, Ben, Cole Trapnell, Mihai Pop, and Steven L. Salzberg. 2009. “Ultrafast and Memory-Efficient Alignment of Short DNA Sequences to the Human Genome.” Genome Biology 10 (3): R25.

39. Liang, Lixiong, Yingying Chang, Junqian Lu, Xiaojuan Wu, Qi Liu, Weixi Zhang, Xiaohua Su, and Bingyu Zhang. 2019. “Global Methylomic and Transcriptomic Analyses Reveal the Broad Participation of DNA Methylation in Daily Gene Expression Regulation of Populus Trichocarpa.” Frontiers in Plant Science 10 (February): 243.

40. Li, Heng. 2018. “Minimap2: Pairwise Alignment for Nucleotide Sequences.” Bioinformatics 34 (18): 3094–3100.

41. Li, Xiang, Kewei Cai, Zhiming Han, Shikai Zhang, Anran Sun, Ying Xie, Rui Han, et al. 2022. “Chromosome-Level Genome Assembly for Acer Pseudosieboldianum and Highlights to Mechanisms for Leaf Color and Shape Change.” Frontiers in Plant Science 13 (March): 850054.

42. Liu, S., Yeh, C.-T., Ji, T., Ying, K., Wu, H., Tang, H. M., Fu, Y., Nettleton, D., & Schnable, P. S. (2009). Mu transposon insertion sites and meiotic recombination events co-localize with epigenetic marks for open chromatin across the maize genome. PLoS Genetics, 5(11), e1000733.

43. Liu, Yang, Wojciech Rosikiewicz, Ziwei Pan, Nathaniel Jillette, Ping Wang, Aziz Taghbalout, Jonathan Foox, et al. 2021. “DNA Methylation-Calling Tools for Oxford Nanopore Sequencing: A Survey and Human Epigenome-Wide Evaluation.” Genome Biology 22 (1): 295.

44. Loman, Nicholas J., Joshua Quick, and Jared T. Simpson. 2015. “A Complete Bacterial Genome Assembled de Novo Using Only Nanopore Sequencing Data.” Nature Methods 12 (8): 733–35.

45. Lu, X., Chen, Z., Liao, B., Han, G., Shi, D., Li, Q., Ma, Q., Zhu, L., Zhu, Z., Luo, X., Fu, S., & Ren, J. (2022). The chromosome-scale genome provides insights into pigmentation in Acer rubrum. Plant Physiology and Biochemistry: PPB / Societe Francaise de Physiologie Vegetale, 186, 322–333.

46. Ma, Qiuyue, Tianlin Sun, Shushun Li, Jing Wen, Lu Zhu, Tongming Yin, Kunyuan Yan, et al. 2020. “The Acer Truncatum Genome Provides Insights into Nervonic Acid Biosynthesis.” The Plant Journal: For Cell and Molecular Biology 104 (3): 662–78.

47. Mao, D., Tao, S., Li, X., Gao, D., Tang, M., Liu, C., Wu, D., Bai, L., He, Z., Wang, X., Yang, L., Zhu, Y., Zhang, D., Zhang, W., & Chen, C. (2022). The Harbinger transposon-derived gene PANDA epigenetically coordinates panicle number and grain size in rice. Plant Biotechnology Journal, 20(6), 1154–1166.

48. Martin, G. T., Seymour, D. K., & Gaut, B. S. (2021). CHH Methylation Islands: A Nonconserved Feature of Grass Genomes That Is Positively Associated with Transposable Elements but Negatively Associated with Gene-Body Methylation. Genome Biology and Evolution, 13(8).

49. McEvoy, Susan L., U. Uzay Sezen, Alexander Trouern-Trend, Sean M. McMahon, Paul G. Schaberg, Jie Yang, Jill L. Wegrzyn, and Nathan G. Swenson. 2021. “Strategies of Tolerance Reflected in Two North American Maple Genomes.” *The Plant Journal: For Cell and Molecular Biology*, December.

50. Mhiri, Corinne, Filipe Borges, and Marie-Angèle Grandbastien. 2022. “Specificities and Dynamics of Transposable Elements in Land Plants.” Biology 11 (4).

51. Miryeganeh, Matin, Ferdinand Marlétaz, Daria Gavriouchkina, and Hidetoshi Saze. 2022. “De Novo Genome Assembly and in Natura Epigenomics Reveal Salinity-Induced DNA Methylation in the Mangrove Tree Bruguiera Gymnorhiza.” The New Phytologist 233 (5): 2094–2110.

52. Moisy, Cédric, Keith E. Garrison, Carole P. Meredith, and Frédérique Pelsy. 2008. “Characterization of Ten Novel Ty1/copia-like Retrotransposon Families of the Grapevine Genome.” BMC Genomics 9 (October): 469.

53. Niederhuth, Chad E., Adam J. Bewick, Lexiang Ji, Magdy S. Alabady, Kyung Do Kim, Qing Li, Nicholas A. Rohr, et al. 2016. “Widespread Natural Variation of DNA Methylation within Angiosperms.” Genome Biology 17 (1): 194.

54. Ni, Peng, Neng Huang, Fan Nie, Jun Zhang, Zhi Zhang, Bo Wu, Lu Bai, et al. 2021. “Genome-Wide Detection of Cytosine Methylations in Plant from Nanopore Sequencing Data Using Deep Learning.” bioRxiv.

55. Ni, Peng, Neng Huang, Zhi Zhang, De-Peng Wang, Fan Liang, Yu Miao, Chuan-Le Xiao, Feng Luo, and Jianxin Wang. 2019. “DeepSignal: Detecting DNA Methylation State from Nanopore Sequencing Reads Using Deep-Learning.” Bioinformatics 35 (22): 4586–95.

56. O’Leary, Nuala A., Mathew W. Wright, J. Rodney Brister, Stacy Ciufo, Diana Haddad, Rich McVeigh, Bhanu Rajput, et al. 2016. “Reference Sequence (RefSeq) Database at NCBI: Current Status, Taxonomic Expansion, and Functional Annotation.” Nucleic Acids Research 44 (D1): D733–45.

57. Orozco-Arias, S., Dupeyron, M., Gutiérrez-Duque, D., Tabares-Soto, R., & Guyot, R. (2023). High nucleotide similarity of three Copia lineage LTR retrotransposons among plant genomes. Genome / National Research Council Canada = Genome / Conseil National de Recherches Canada, 66(3), 51–61.

58. Orozco-Arias, S., Jaimes, P. A., Candamil, M. S., Jiménez-Varón, C. F., Tabares-Soto, R., Isaza, G., & Guyot, R. (2021). InpactorDB: A Classified Lineage-Level Plant LTR Retrotransposon Reference Library for Free-Alignment Methods Based on Machine Learning. Genes, 12(2).

59. Oxford Nanopore Technologies. 2020a. *Guppy*. Github. https://github.com/nanoporetech. 2020b. *Tombo* (version 1.5.1). https://github.com/nanoporetech/tombo. 2021. *ont_fast5_api* (version 4.0.0). https://github.com/nanoporetech/ont_fast5_api.

60. Perumal, Sampath, Chu Shin Koh, Lingling Jin, Miles Buchwaldt, Erin E. Higgins, Chunfang Zheng, David Sankoff, et al. 2020. “A High-Contiguity Brassica Nigra Genome Localizes Active Centromeres and Defines the Ancestral Brassica Genome.” Nature Plants 6 (8): 929–41.

61. Pratama, Sigit Nur, Fenny Martha Dwivany, and Husna Nugrahapraja. 2021. “Comparative Genomics of Copia and Gypsy Retroelements in Three Banana Genomes: A, B, and S Genomes.” Pertanika Journal of Tropical Agricultural Science 44 (4).

62. Qiu, Fan, and Mark C. Ungerer. 2018. “Genomic Abundance and Transcriptional Activity of Diverse Gypsy and Copia Long Terminal Repeat Retrotransposons in Three Wild Sunflower Species.” BMC Plant Biology 18 (1): 6.

63. Quinlan, Aaron R., and Ira M. Hall. 2010. “BEDTools: A Flexible Suite of Utilities for Comparing Genomic Features.” Bioinformatics 26 (6): 841–42.

64. R Core Team. 2013. “R: A Language and Environment for Statistical Computing.” Vienna, Austria: R Foundation for Statistical Computing. http://www.R-project.org/.

65. Ritter, Eleanore J., and Chad E. Niederhuth. 2021. “Intertwined Evolution of Plant Epigenomes and Genomes.” Current Opinion in Plant Biology 61 (June): 101990.

66. Rose, Alan B. 2018. “Introns as Gene Regulators: A Brick on the Accelerator.” Frontiers in Genetics 9: 672.

67. Ruggieri, V., Alexiou, K. G., Morata, J., Argyris, J., Pujol, M., Yano, R., Nonaka, S., Ezura, H., Latrasse, D., Boualem, A., Benhamed, M., Bendahmane, A., Cigliano, R. A., Sanseverino, W., Puigdomènech, P., Casacuberta, J. M., & Garcia-Mas, J. (2018). An improved assembly and annotation of the melon (Cucumis melo L.) reference genome. Scientific Reports, 8(1), 8088.

68. Sasaki, Eriko, Taiji Kawakatsu, Joseph R. Ecker, and Magnus Nordborg. 2019. “Common Alleles of CMT2 and NRPE1 Are Major Determinants of CHH Methylation Variation in Arabidopsis Thaliana.” PLoS Genetics 15 (12): e1008492.

69. Shi, Danlu, Kai Zhuang, Yan Xia, Changhua Zhu, Chen Chen, Zhubing Hu, and Zhenguo Shen. 2017. “Hydrilla Verticillata Employs Two Different Ways to Affect DNA Methylation under Excess Copper Stress.” Aquatic Toxicology 193 (December): 97–104.

70. Smit, A. F. A., R. Hubley, and P. Green. 2013-2015. RepeatMasker Open*-*4.0. http://www.repeatmasker.org.

71. Sork, Victoria L., Shawn J. Cokus, Sorel T. Fitz-Gibbon, Aleksey V. Zimin, Daniela Puiu, Jesse A. Garcia, Paul F. Gugger, et al. 2022. “High-Quality Genome and Methylomes Illustrate Features Underlying Evolutionary Success of Oaks.” Nature Communications 13 (1): 2047.

72. Stritt, C., Wyler, M., Gimmi, E. L., Pippel, M., & Roulin, A. C. (2020). Diversity, dynamics and effects of long terminal repeat retrotransposons in the model grass Brachypodium distachyon. The New Phytologist, 227(6), 1736–1748.

73. Suguiyama, V. F., Vasconcelos, L. A. B., Rossi, M. M., Biondo, C., & de Setta, N. (2019). The population genetic structure approach adds new insights into the evolution of plant LTR retrotransposon lineages. PloS One, 14(5), e0214542.

74. Underwood, C. J., & Choi, K. (2019). Heterogeneous transposable elements as silencers, enhancers and targets of meiotic recombination. Chromosoma, 128(3), 279–296.

75. UniProt Consortium. 2019. “UniProt: A Worldwide Hub of Protein Knowledge.” Nucleic Acids Research 47 (D1): D506–15.

76. Vaillancourt, Brieanne, and C. Robin Buell. 2019. “High Molecular Weight DNA Isolation Method from Diverse Plant Species for Use with Oxford Nanopore Sequencing.” bioRxiv.

77. Vitte, C., Fustier, M.-A., Alix, K., & Tenaillon, M. I. (2014). The bright side of transposons in crop evolution. Briefings in Functional Genomics, 13(4), 276–295.

78. Walker, Bruce J., Thomas Abeel, Terrance Shea, Margaret Priest, Amr Abouelliel, Sharadha Sakthikumar, Christina A. Cuomo, et al. 2014. “Pilon: An Integrated Tool for Comprehensive Microbial Variant Detection and Genome Assembly Improvement.” PloS One 9 (11): e112963.

79. Wang, Y., Liang, W., & Tang, T. (2018). Constant conflict between Gypsy LTR retrotransposons and CHH methylation within a stress-adapted mangrove genome. The New Phytologist, 220(3), 922–935.

80. Wenke, T., Döbel, T., Sörensen, T. R., Junghans, H., Weisshaar, B., & Schmidt, T. (2011). Targeted identification of short interspersed nuclear element families shows their widespread existence and extreme heterogeneity in plant genomes. The Plant Cell, 23(9), 3117–3128.

81. Yang, Jing, Hafiz Muhammad Wariss, Lidan Tao, Rengang Zhang, Quanzheng Yun, Peter Hollingsworth, Zhiling Dao, et al. 2019. “De Novo Genome Assembly of the Endangered Acer Yangbiense, a Plant Species with Extremely Small Populations Endemic to Yunnan Province, China.” GigaScience 8 (7). https://doi.org/10.1093/gigascience/giz085.

82. Yang, Zhaoen, Xiaoyang Ge, Weinan Li, Yuying Jin, Lisen Liu, Wei Hu, Fuyan Liu, Yanli Chen, Shaoliang Peng, and Fuguang Li. 2021. “Cotton D Genome Assemblies Built with Long-Read Data Unveil Mechanisms of Centromere Evolution and Stress Tolerance Divergence.” BMC Biology 19 (1): 115.

83. Yuen, Zaka Wing-Sze, Akanksha Srivastava, Runa Daniel, Dennis McNevin, Cameron Jack, and Eduardo Eyras. 2021. “Systematic Benchmarking of Tools for CpG Methylation Detection from Nanopore Sequencing.” Nature Communications 12 (1): 3438.

84. Yu, Tao, Yiheng Hu, Yuyang Zhang, Ran Zhao, Xueqing Yan, Buddhi Dayananda, Jinpeng Wang, Yuannian Jiao, Junqing Li, and Xin Yi. 2021. “Whole-Genome Sequencing of Acer Catalpifolium Reveals Evolutionary History of Endangered Species.” Genome Biology and Evolution 13 (12).

85. Zagorski, Danijela, Matthias Hartmann, Yann J. K. Bertrand, Ladislava Paštová, Renata Slavíková, Jiřina Josefiová, and Judith Fehrer. 2020. “Characterization and Dynamics of Repeatomes in Closely Related Species of Hieracium (Asteraceae) and Their Synthetic and Apomictic Hybrids.” Frontiers in Plant Science 11 (November): 591053.

86. Zakrzewski, F., Schmidt, M., Van Lijsebettens, M., & Schmidt, T. (2017). DNA methylation of retrotransposons, DNA transposons and genes in sugar beet (Beta vulgaris L.). The Plant Journal: For Cell and Molecular Biology, 90(6), 1156–1175.

87. Zhang, X., Jiang, N., Feschotte, C., & Wessler, S. R. (2004). PIF-and Pong-Like Transposable Elements: Distribution, Evolution and Relationship With Tourist-Like Miniature Inverted-Repeat Transposable Elements. Genetics, 166(2), 971–986.

88. Zimin, Aleksey V., Daniela Puiu, Ming-Cheng Luo, Tingting Zhu, Sergey Koren, Guillaume Marçais, James A. Yorke, Jan Dvořák, and Steven L. Salzberg. 2017. “Hybrid Assembly of the Large and Highly Repetitive Genome of Aegilops Tauschii, a Progenitor of Bread Wheat, with the MaSuRCA Mega-Reads Algorithm.” Genome Research 27 (5): 787–92.

