## Supplementary figures and images for "Profiling genome-wide methylation in two maples: fine-scale approaches to detection with nanopore technology"

### Supplemental Figure 1

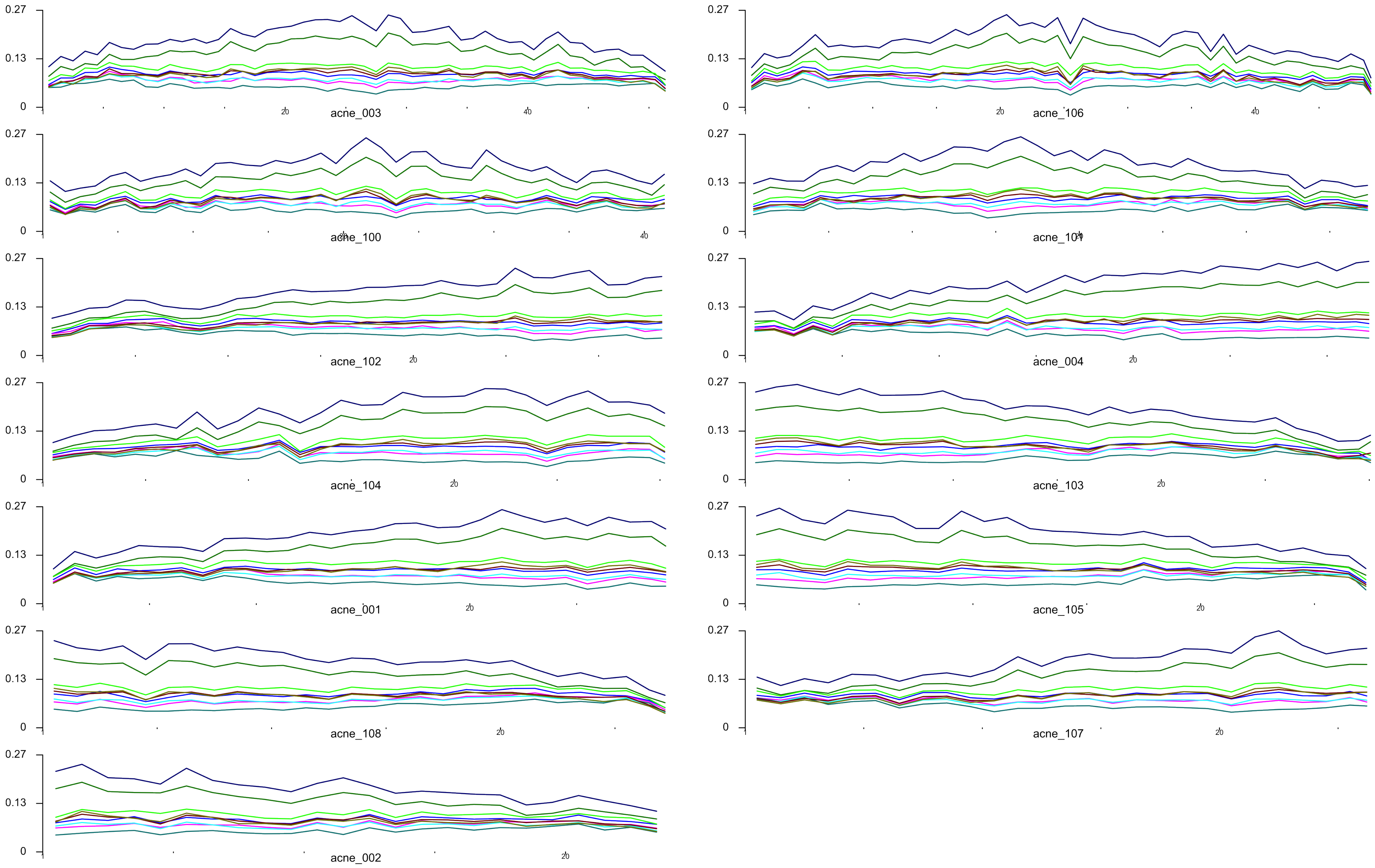

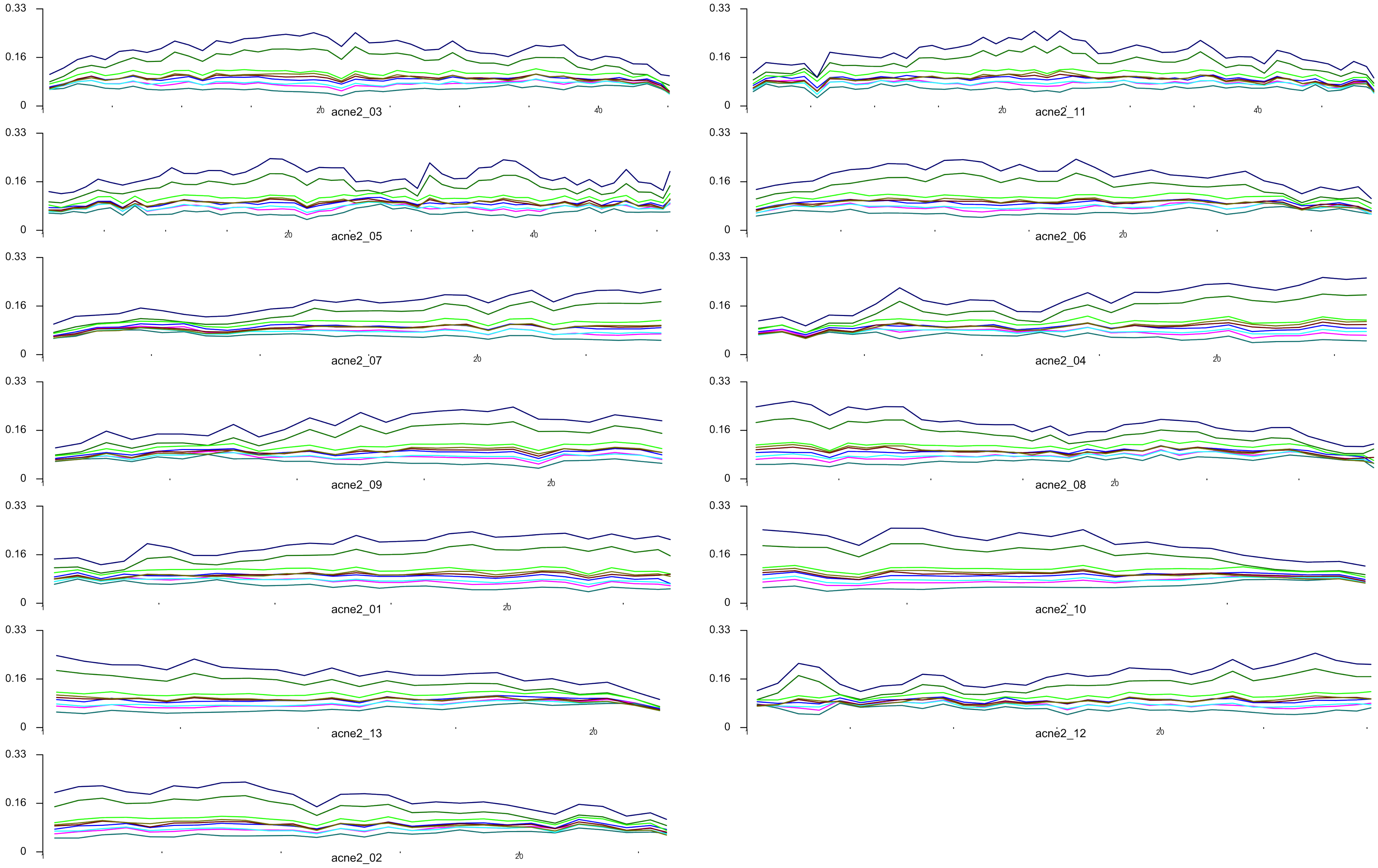

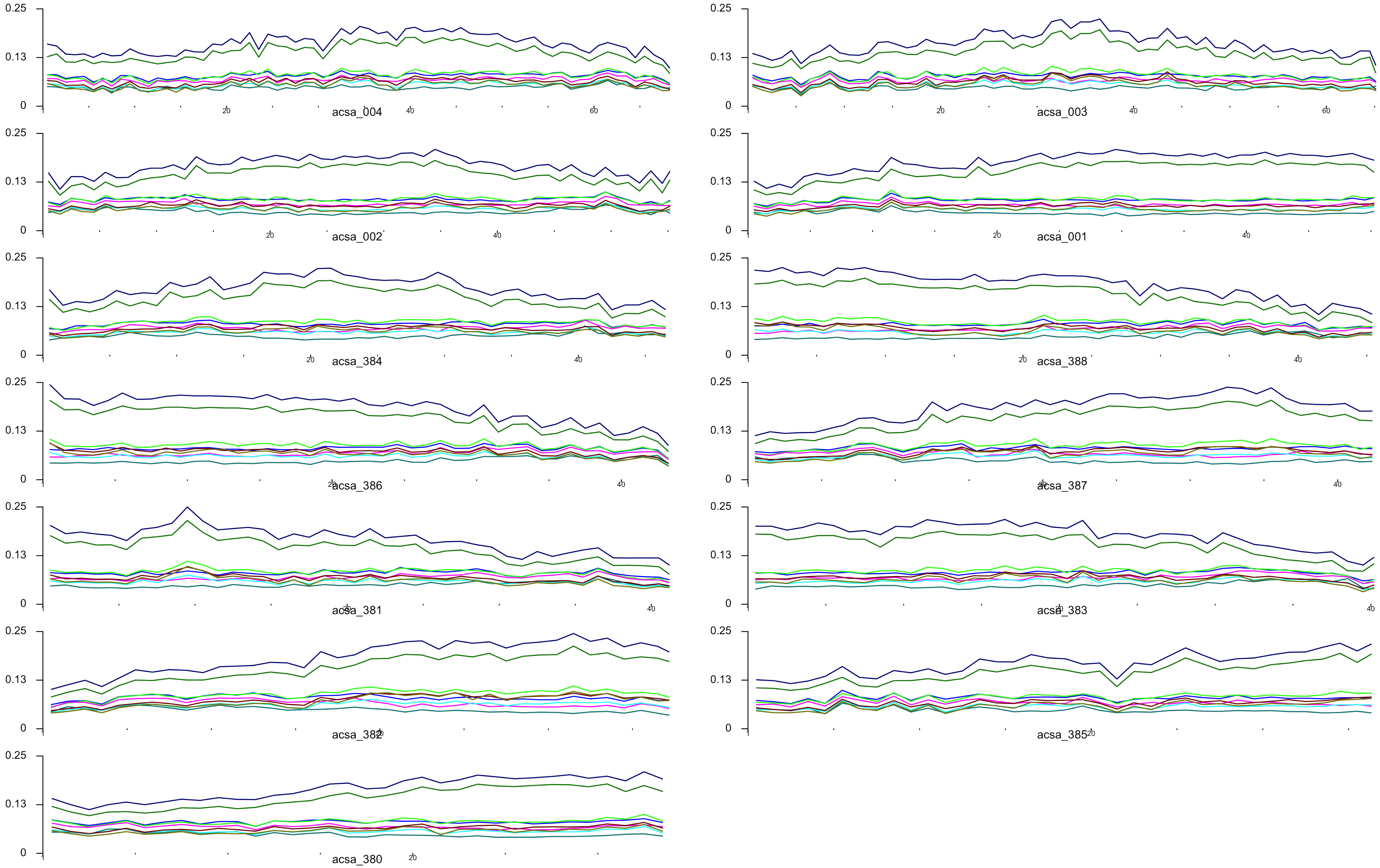

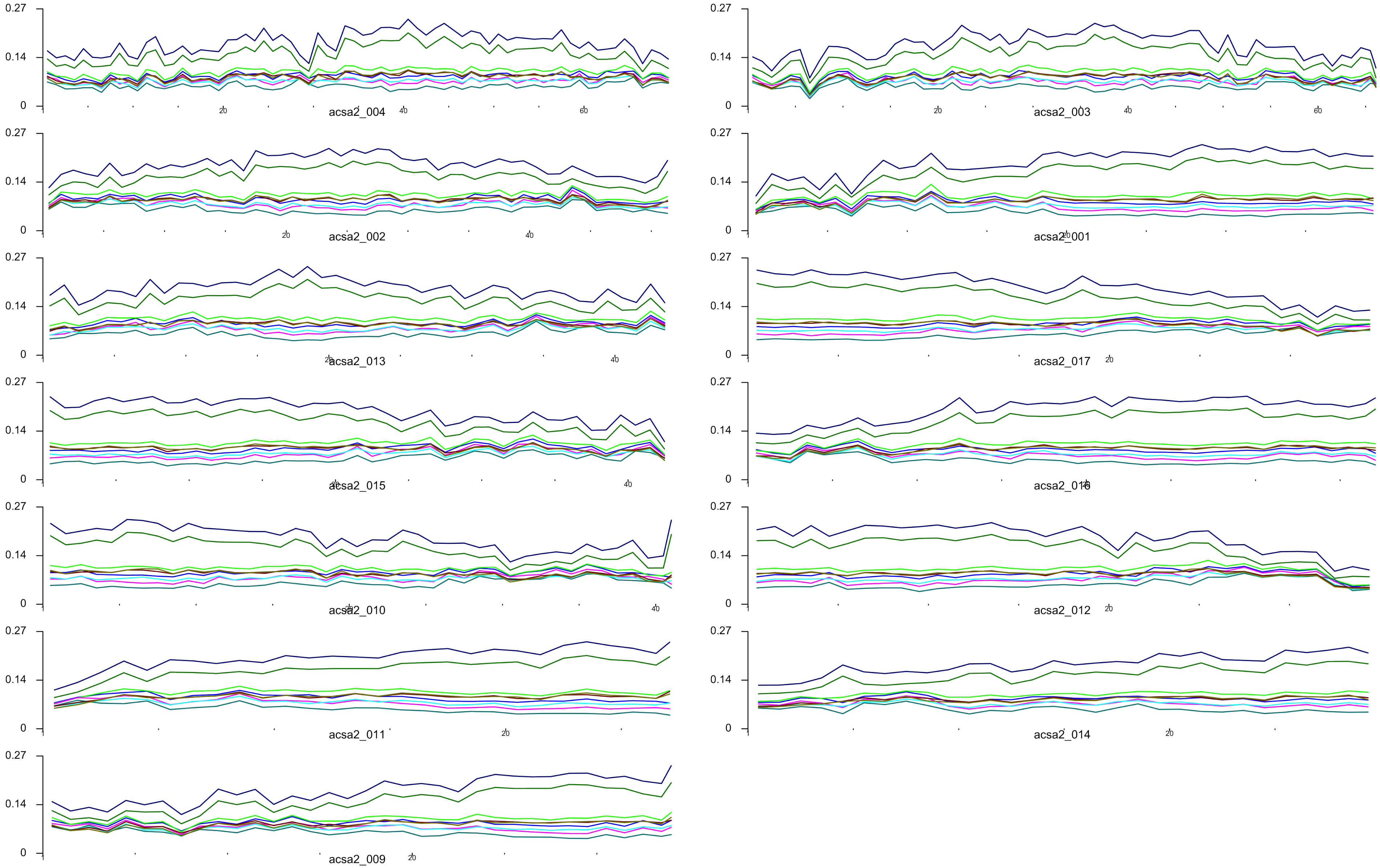

### Supplemental Figure 2

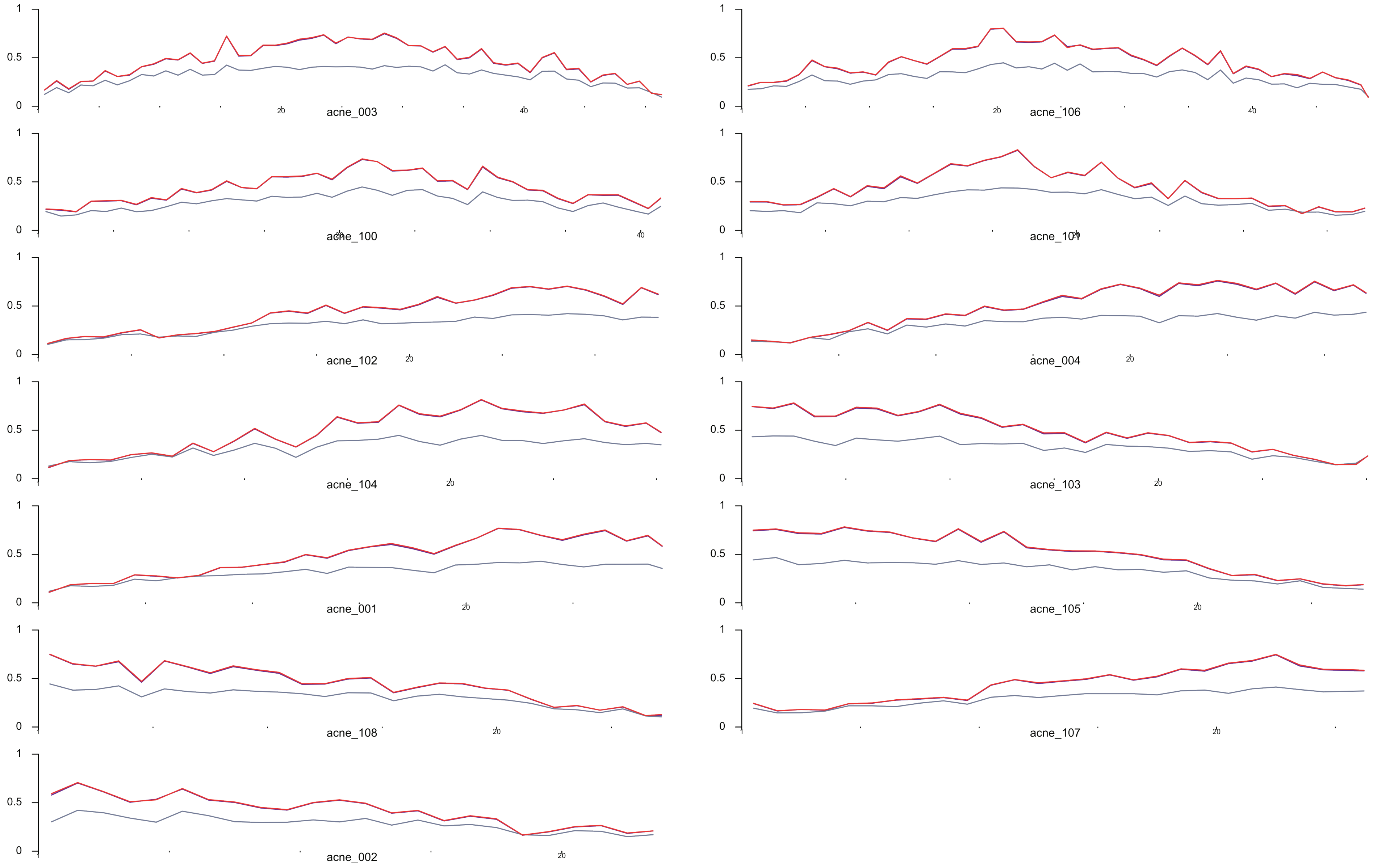

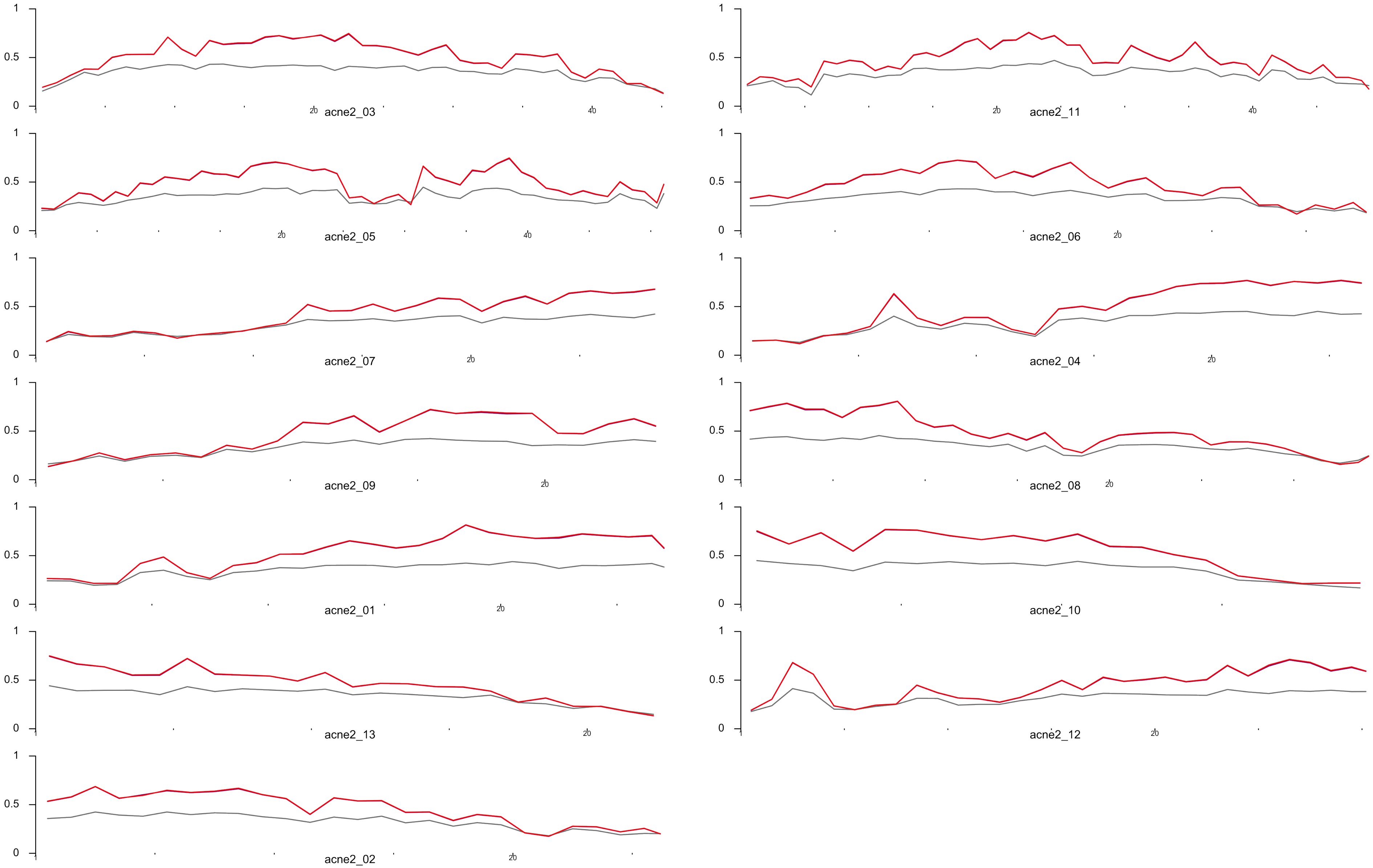

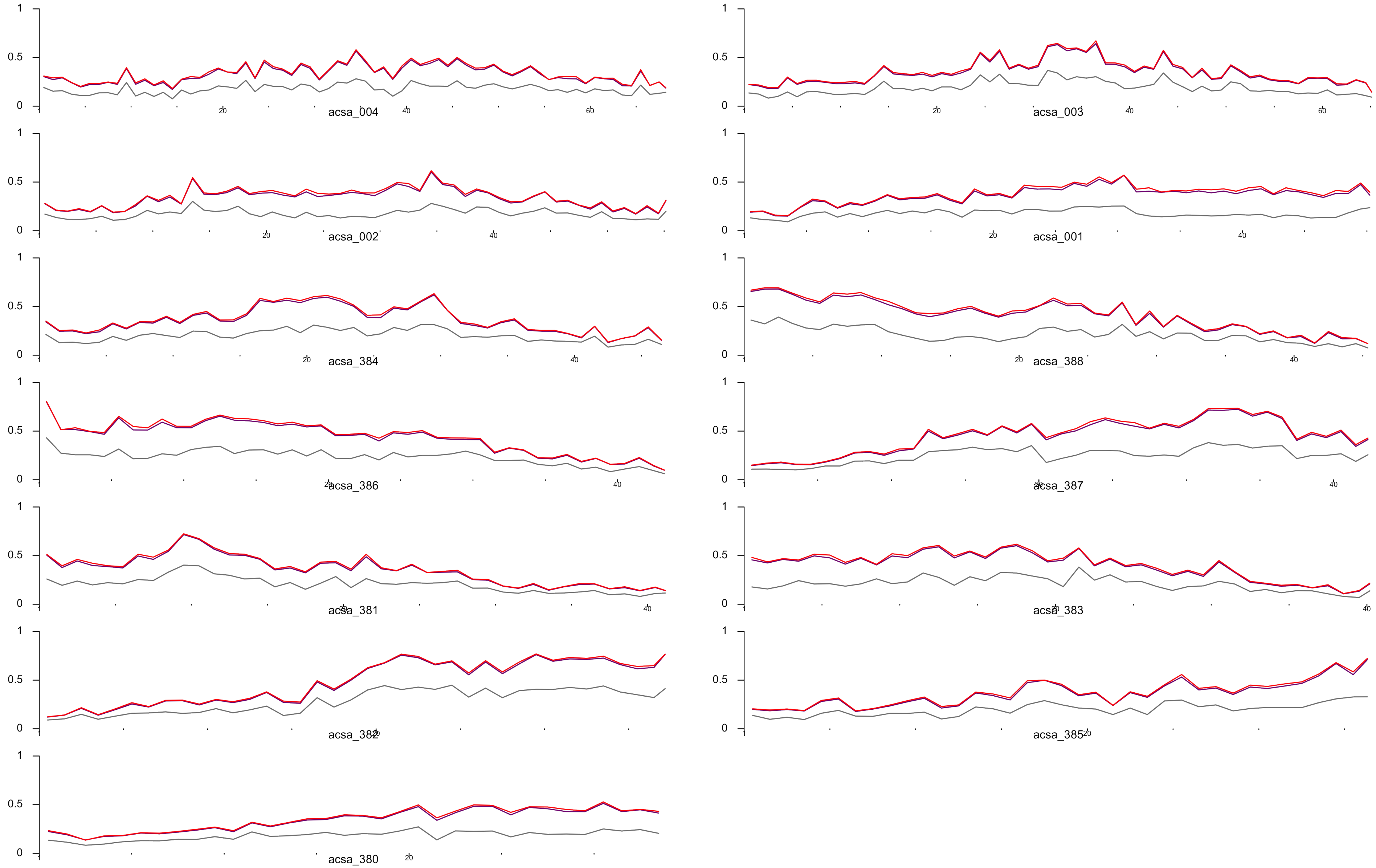

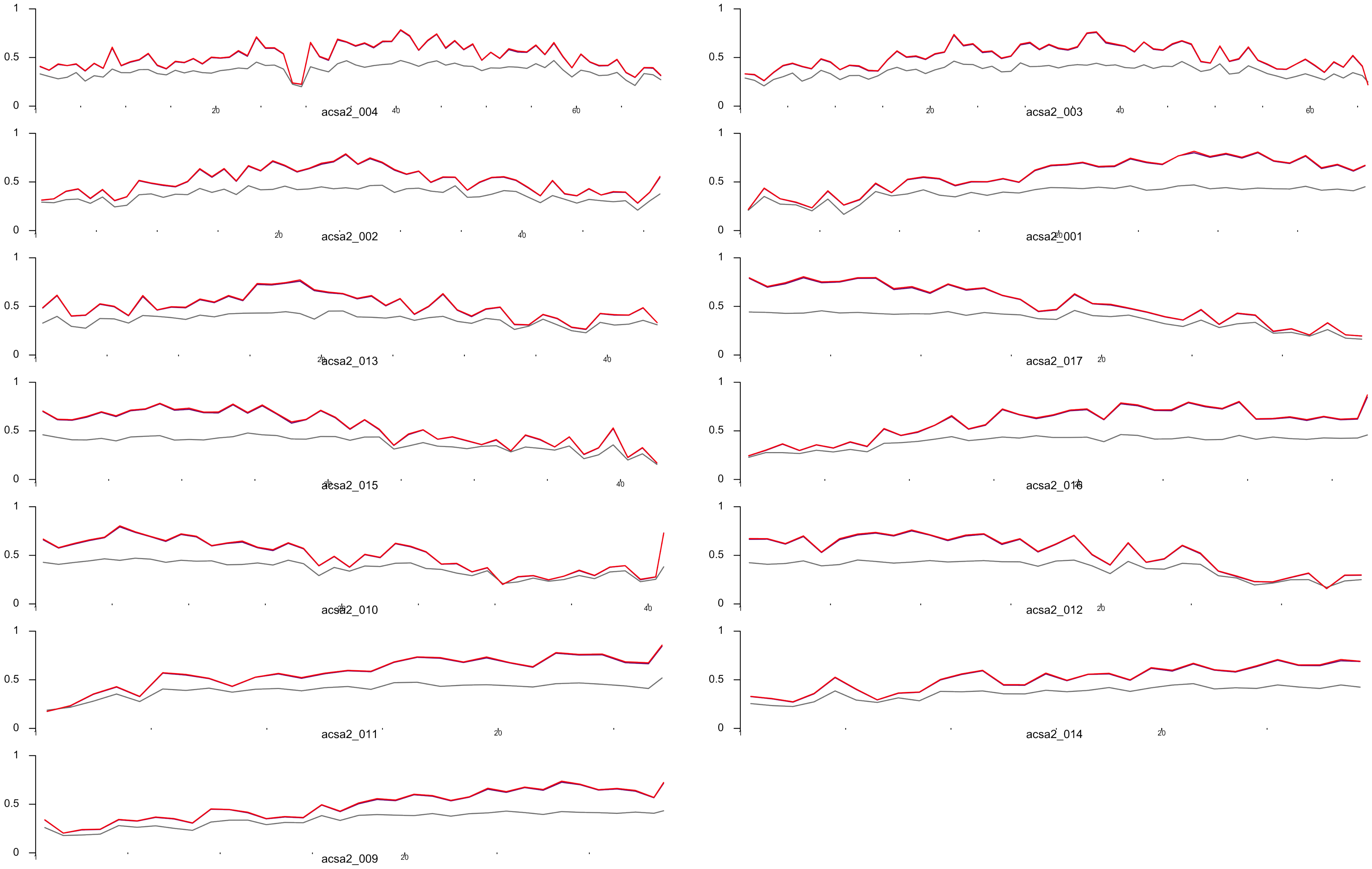

### Supplemental Figure 3

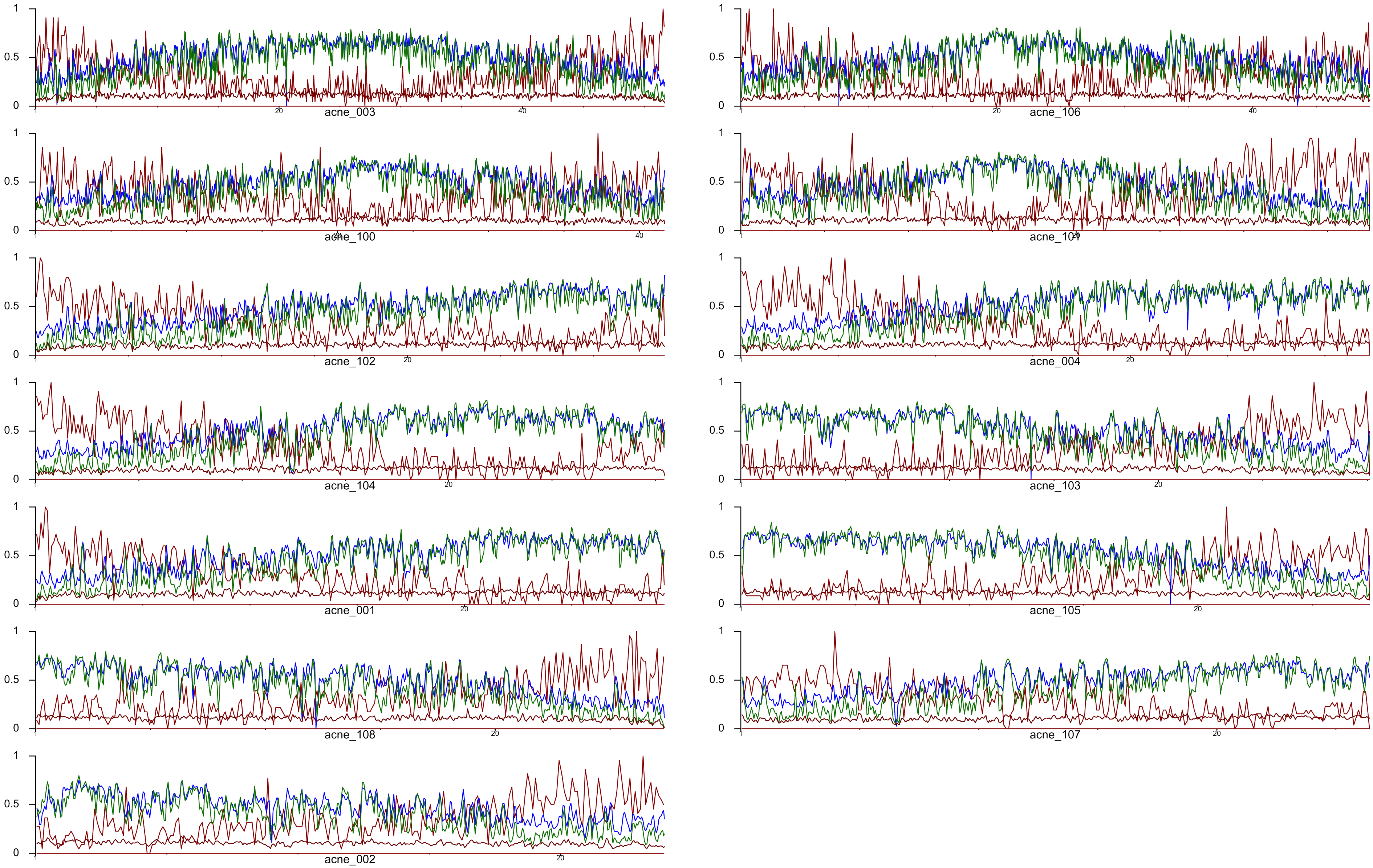

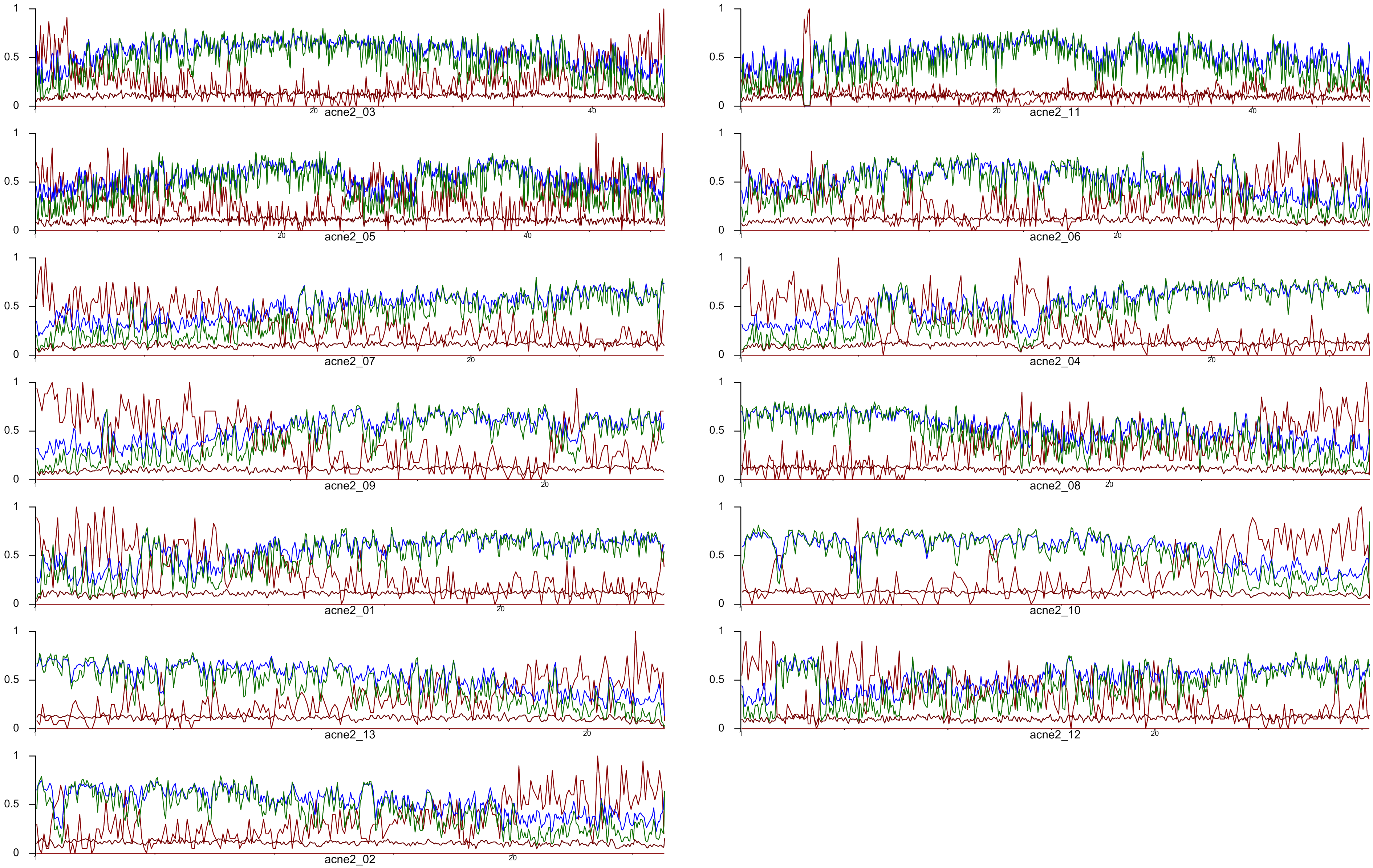

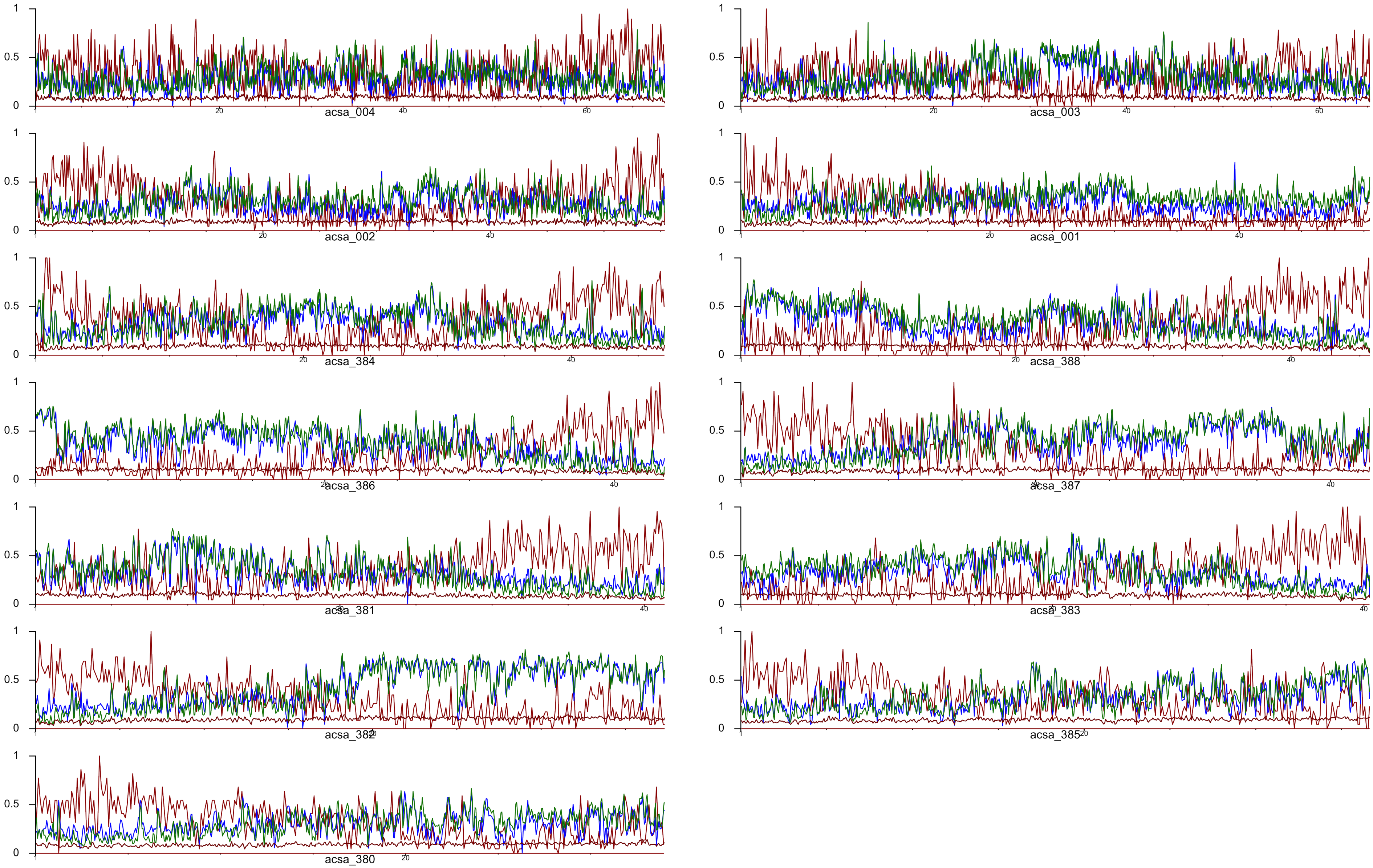

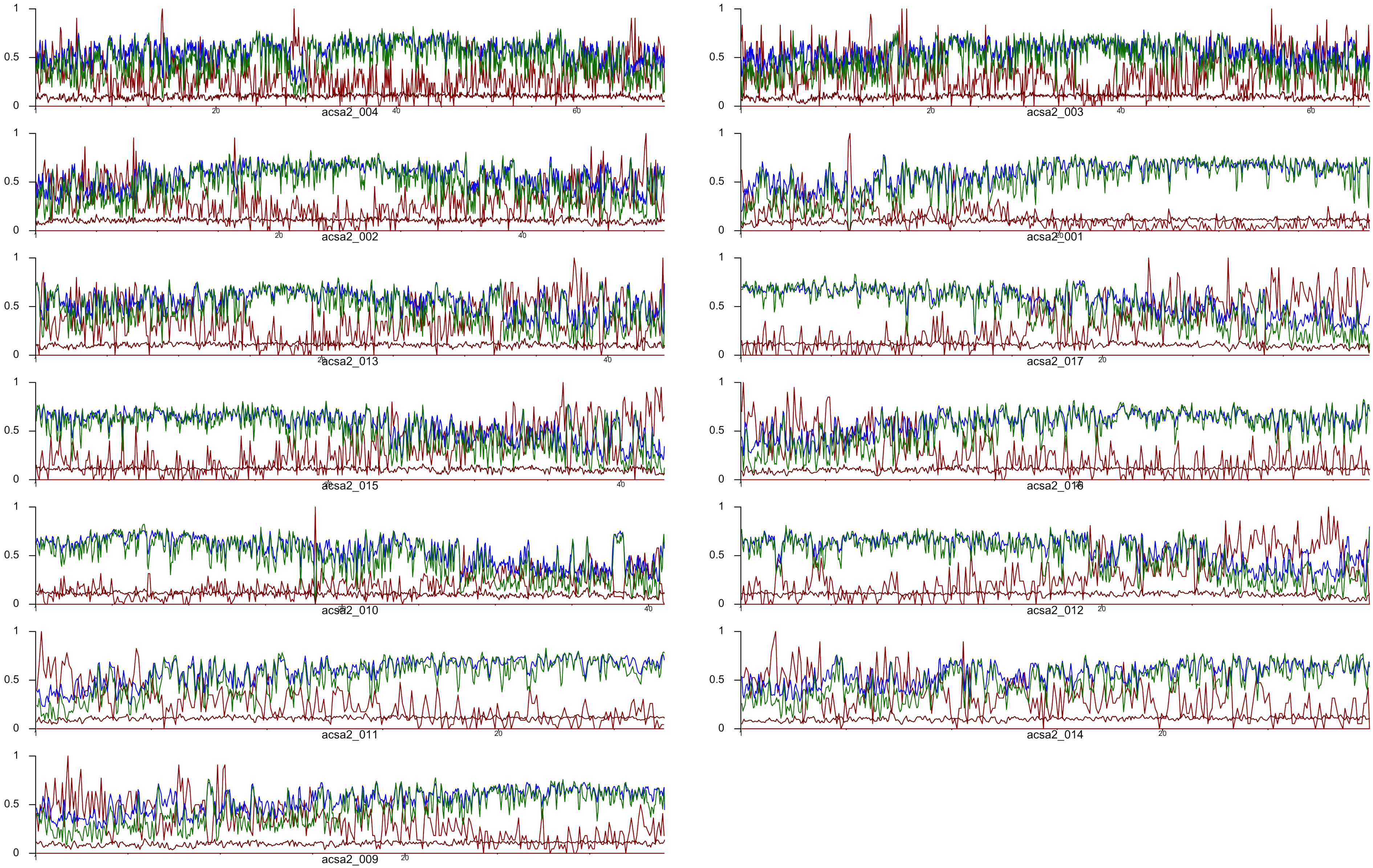

### Supplemental Figure 4

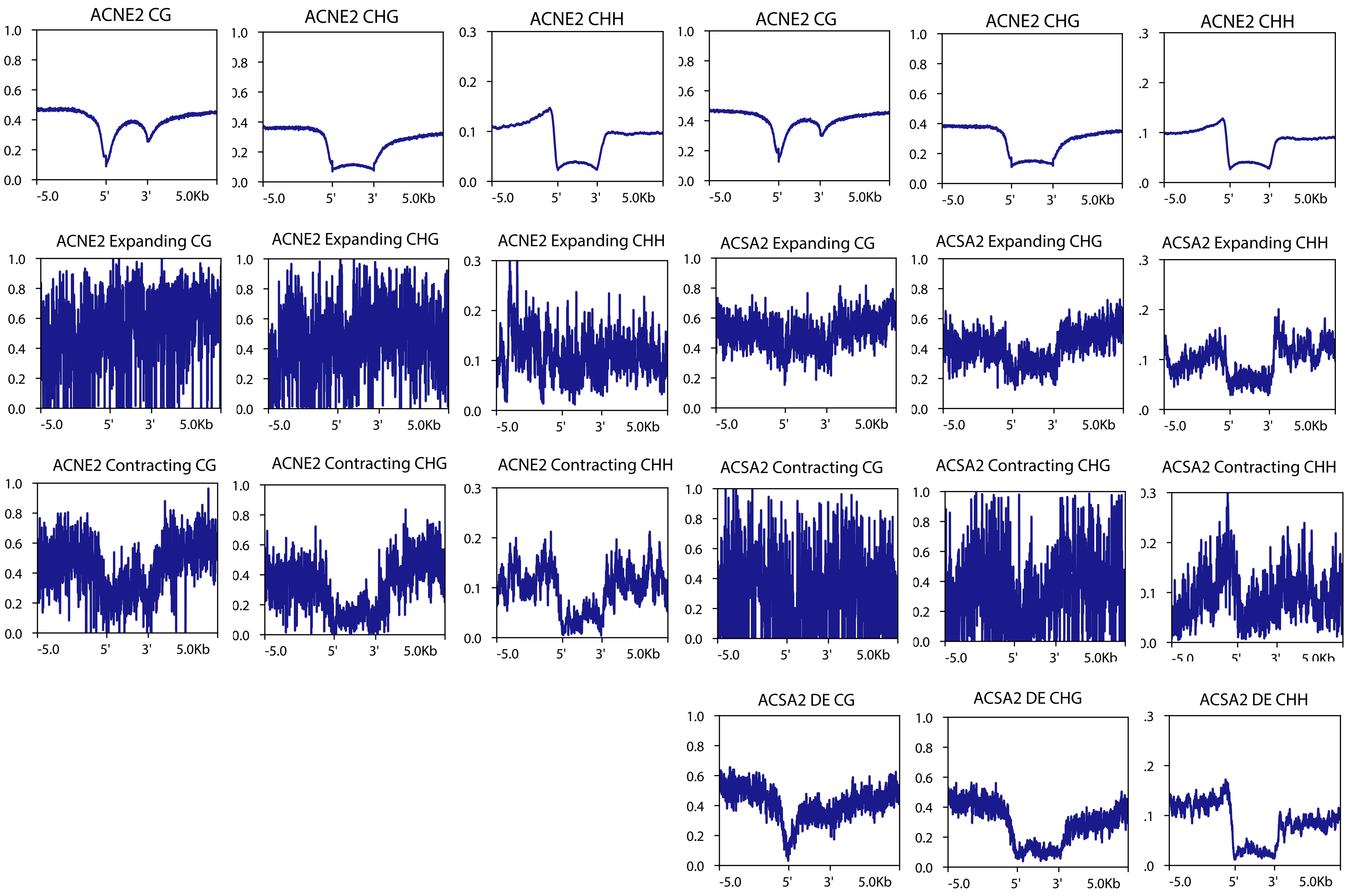

### Supplemental Figure 5

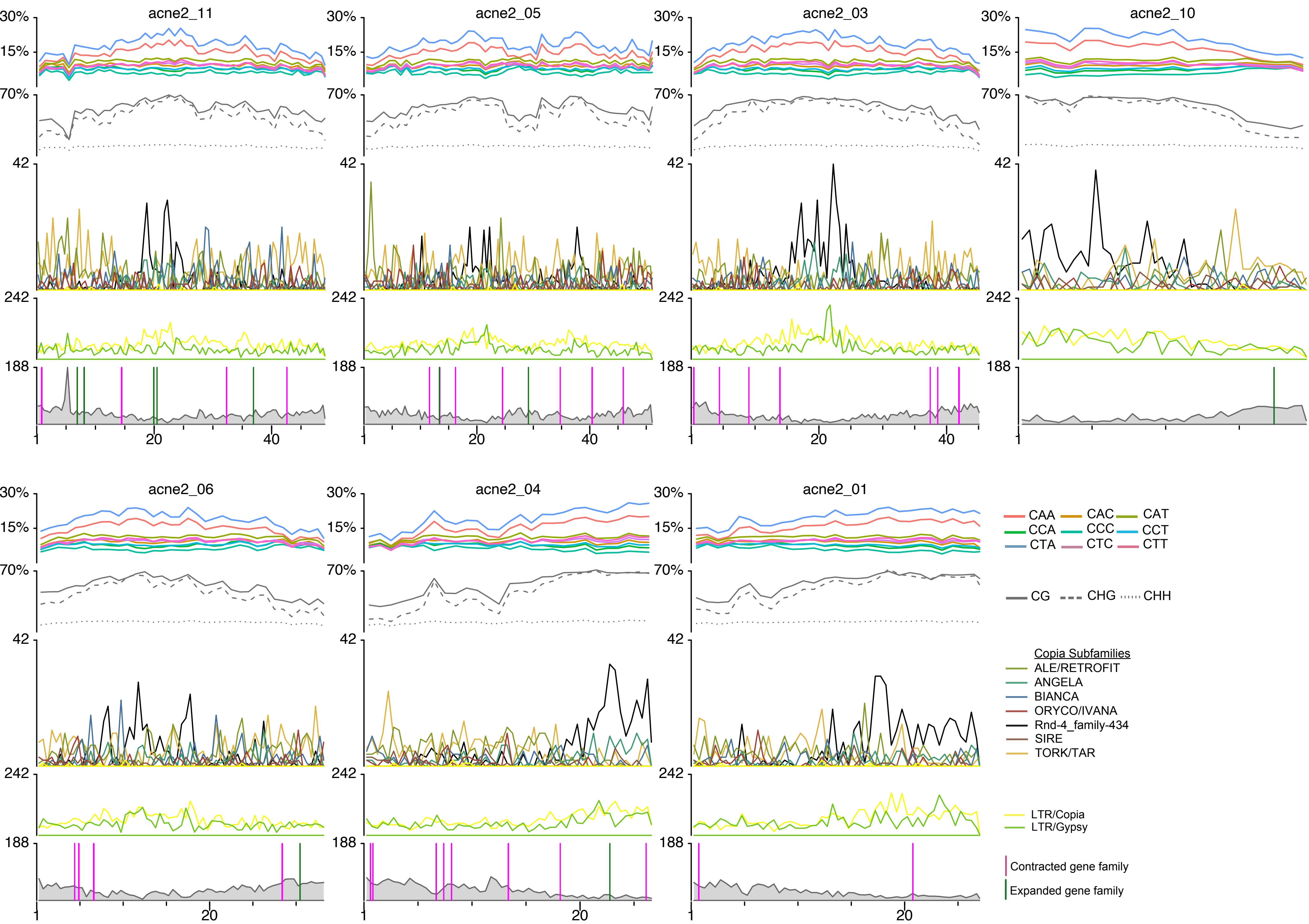

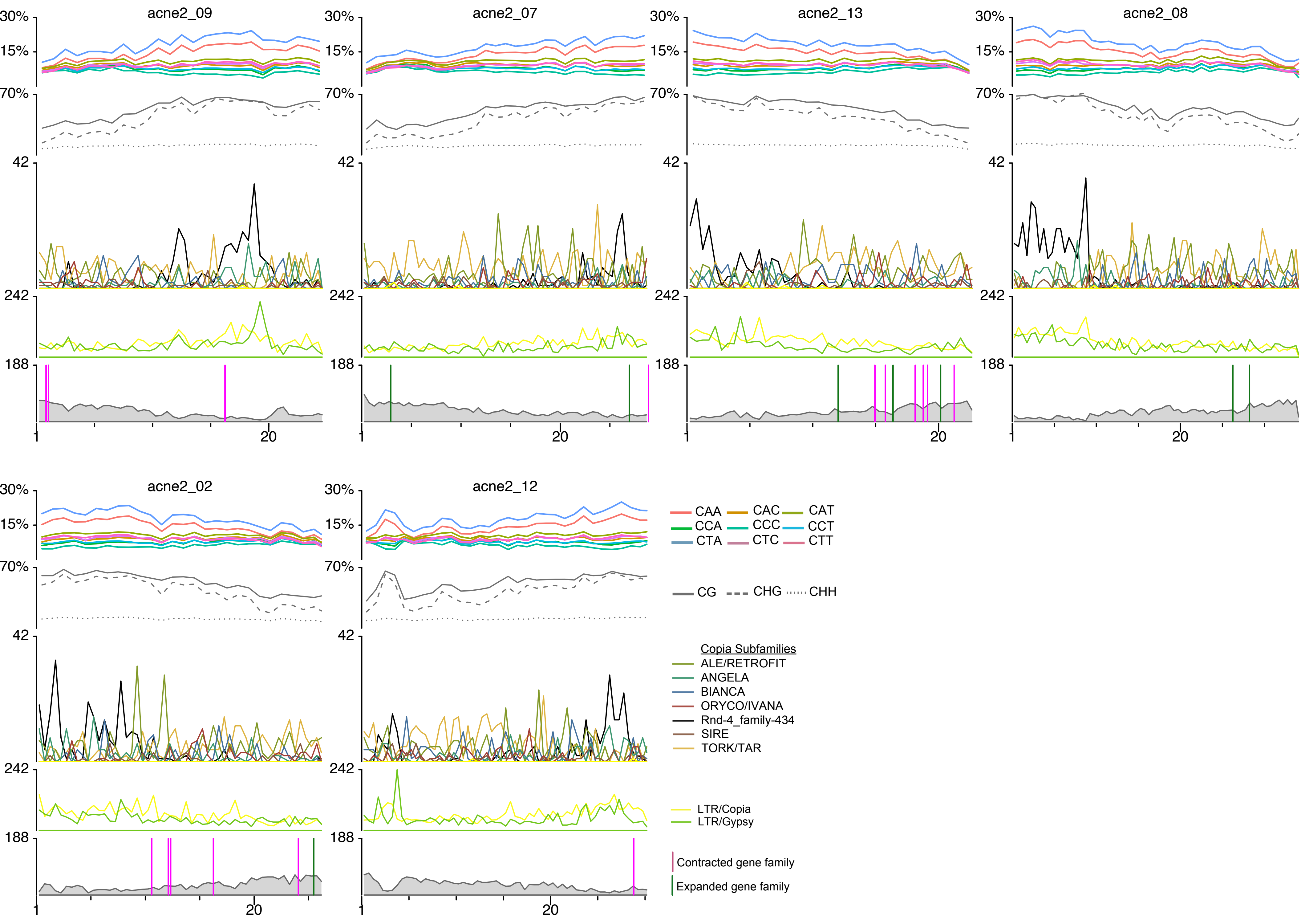

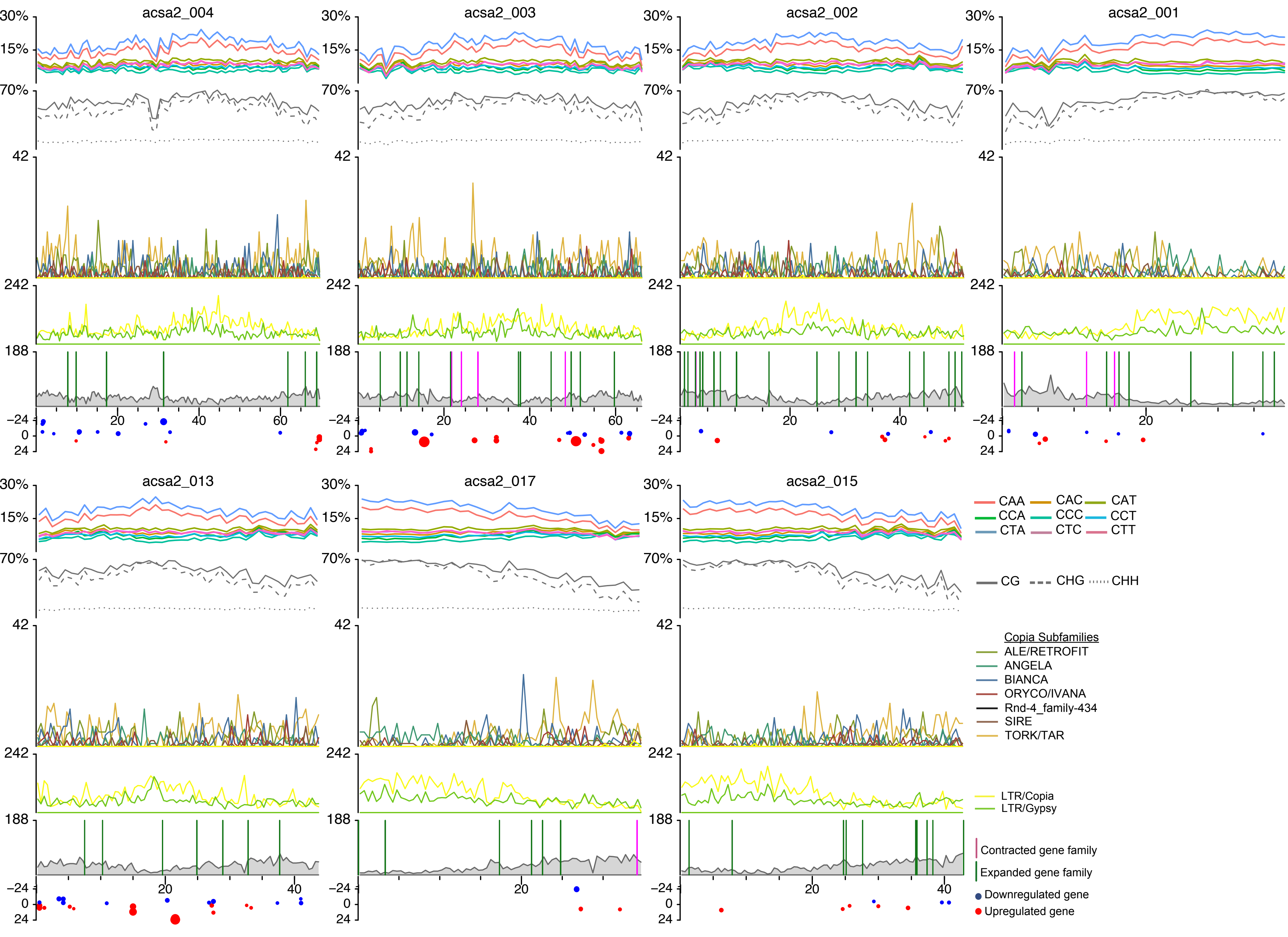

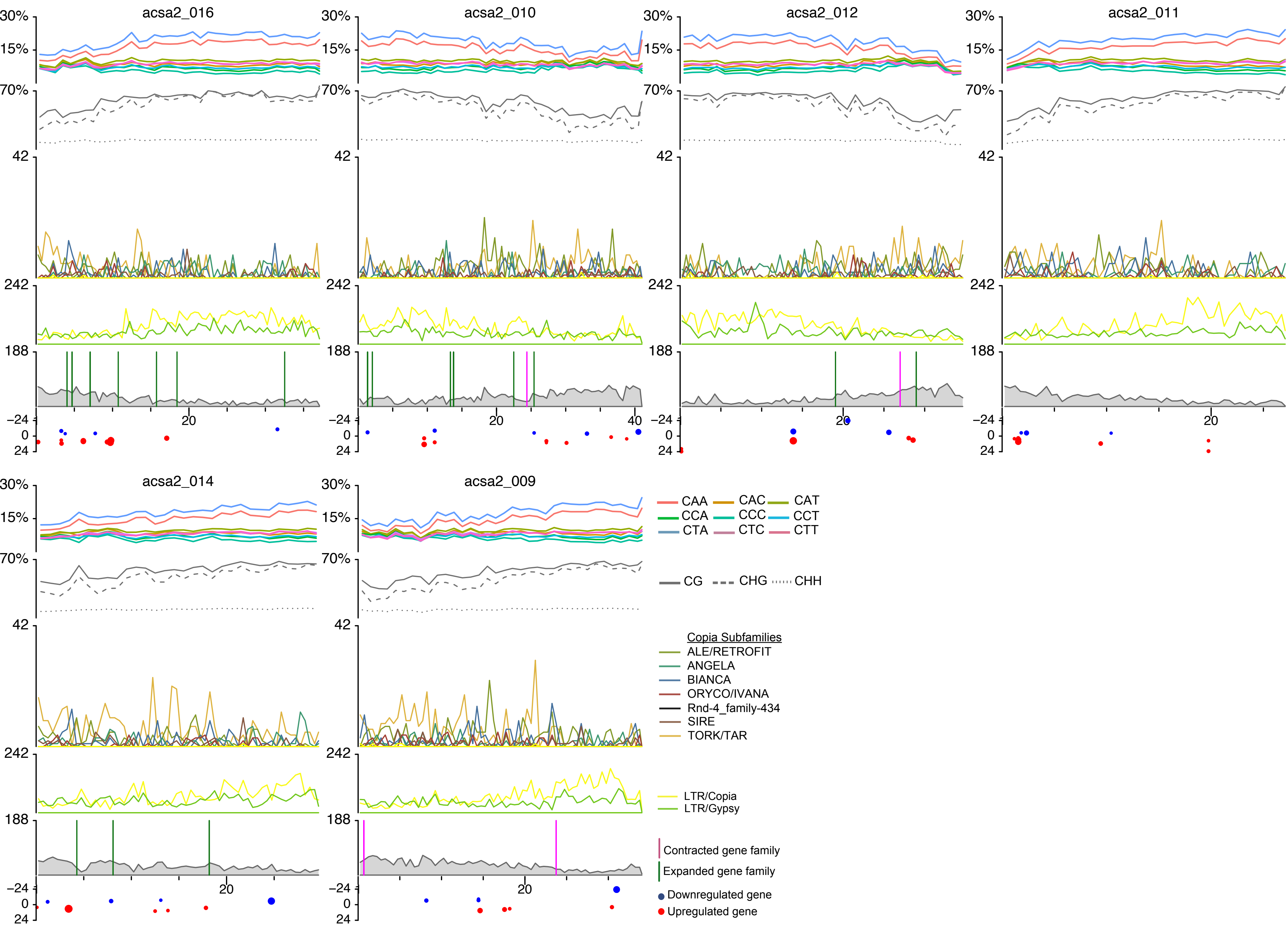
